# Single-Cell Analysis Identifies Thymic Maturation Delay in Growth-Restricted Neonatal Mice

**DOI:** 10.1101/372862

**Authors:** Wendi A. Bacon, Russell S. Hamilton, Ziyi Yu, Jens Kieckbusch, Delia Hawkes, Ada M. Krzak, Chris Abell, Francesco Colucci, D. Stephen Charnock-Jones

## Abstract

Fetal growth restriction (FGR) causes a wide variety of defects in the neonate which can lead to increased risk of heart disease, diabetes, anxiety and other disorders later in life. However, the effect of FGR on the immune system, is poorly understood. We used a well-characterized mouse model of FGR in which placental Igf-2 production is lost due to deletion of the placental specific *Igf-2* P_0_ promotor. The thymi in such animals were reduced in mass with a ∼70% reduction in cellularity. We used single cell RNA sequencing (Drop-Seq) to analyze 7264 thymus cells collected at postnatal day 6.

We identified considerable heterogeneity among the Cd8/Cd4 double positive cells with one subcluster showing marked upregulation of transcripts encoding a sub-set of proteins that contribute to the surface of the ribosome. The cells from the FGR animals were underrepresented in this cluster. Furthermore, the distribution of cells from the FGR animals was skewed with a higher proportion of immature double negative cells and fewer mature T-cells. Cell cycle regulator transcripts also varied across clusters. The T-cell deficit in FGR mice persisted into adulthood, even when body and organ weights approached normal levels due to catch-up growth. This finding complements the altered immunity found in growth restricted human infants. This reduction in T-cellularity may have implications for adult immunity, adding to the list of adult conditions in which the *in utero* environment is a contributory factor.

## 1 Introduction

Fetal growth restriction (FGR) is a serious condition affecting between 3 and 7% of infants in developed nations (1) but can be higher than 40% in low-income nations (2). The growth restricted infants suffer from an increased risk of a wide variety of acute complications, including necrotizing enterocolitis, perinatal asphyxia, hypothermia, hypoglycemia, immunodeficiency and death (3). FGR is commonly attributed to insufficient fetal nutrients. This may be because either the mother is malnourished, as in the prototypical case of the Dutch Hunger Winter (4), or because the placenta is dysfunctional, leading to poor nutrient transport to the fetus (5). Severely growth restricted infants exhibit asymmetric growth wherein resources are reallocated to the fetal brain at the cost of the rest of the body (6), so called ‘brain sparing’. The infants undergo ‘catch-up growth’ for years, although some may remain small for life (7). Unfortunately, this rapid growth often fails to overcome the complications of intrauterine restriction, as a wide variety of morbidities including diabetes (8), polycystic ovary syndrome (9) or perturbed mental health (10) have been linked to FGR or catch-up growth (11). Together, these support the ‘Developmental Origins of Adult Disease’ paradigm, wherein adverse factors during critical periods of fetal growth irreversibly impact development (11,12). To overcome these effects, a clear understanding of the mechanism for each of these sequelae is necessary (13).

Several animal models have been developed for the study of adult diseases originating in the uterus. Most of these involve restricting certain nutritional components, such as protein (14) or oxygen (15) in pregnant dams. However, this does not distinguish between maternal placental effects. The *Igf-2*^*P0*^model overcomes this through knock-out of placenta-specific insulin-like growth factor 2. There are four *Igf-2* isoforms which result from the use of different promoters although they ultimately generate the same protein. The P_0_ promoter is specific to the placenta. This gene is paternally imprinted, allowing for generation of both wildtype and affected offspring within the same litter. Importantly, all offspring develop in a wildtype dam, preventing maternal variables from affecting development. This targeted knock-out reduces placental growth and therefore the nutrient transport to the fetus, resulting in a brain-sparing phenotype reminiscent of human FGR (16). Early hypocalcemia in the fetuses of these mice (17) mimics the hypocalcemia found in human neonates (18). These mice have been shown to develop anxiety later in life (19), which recapitulates known long-term effects of FGR on mental health (20).

While long-term effects are a subject of much interest, most acute FGR complications are simply attributed to a lack of tissue mass and developmental delay: for example, a smaller and less mature kidney (21), pancreas (22) or bowel (23) will simply not function as well. Adaptive immunity is mediated by T-cells which develop in the thymus. However, while the thymus is a short-lived organ which involutes shortly after birth it continues to function well into adult life (24). Deleterious effects on this transient organ could, therefore, have a significant and irreversible impact on immunity in adult life. Initially, infants with FGR have acutely smaller thymi and altered CD4/CD8 ratios of peripheral T cells (25). Later in life, FGR is associated with abnormal responses to vaccines and higher rates of death due to infection (26). For example, indirect evidence comes from a study showing that young adults born in the annual ‘hungry season’ in rural Gambia – and therefore likely to be born with FGR – have a 10-fold higher risk of premature death, largely due to infection (27).

At a cellular level, broad definitions classify cells based on discrete cell-surface markers. In T-cell development, lymphoid progenitors travel from the bone marrow through the bloodstream to arrive at the thymus where NOTCH signaling directs them towards the T-cell lineage (28). These cells divide and differentiate through four stages of DN (double negative, referring to lack of either CD4 or CD8 T-cell surface markers) whilst undergoing *Tcr*_*β*_ rearrangements in the thymic cortex. With the correct set of signals, a minority of the cells become γδ T cells. The rest gain CD8 and then CD4 surface makers (DP, double-positive) at which point *Tcr*_*α*_ rearrangements take place. The rearrangements provide the diverse T-cell receptor repertoire necessary for antigen recognition. The cell is then activated by either MHC Class I or Class II molecules presenting self-antigens. Strong affinity for either MHC Class I or Class II leads to apoptosis to prevent auto-immune disease. Weak affinity to Class I or II leads to differentiation into either CD8+ cytotoxic T cells or CD4+ T helper cells, respectively. Intermediate affinity to Class II molecules leads to differentiation into a T-regulatory cell. Errors anywhere along this pathway of T-cell differentiation, from insufficient *Tcr* rearrangements to underrepresentation of any of the T cell lineages, can lead to impaired immune function (29), (30).

Single-cell RNA sequencing, for example Drop-Seq (31), In-Drop, or the commercial 10X Genomics and Dolomite platforms allow the analysis of the transcriptomes of thousands of single-cells (32). These analyses have been invaluable for identifying immune-cell subtypes within populations traditionally classified by discrete cell surface markers (33) and revealed new regulatory pathways (34).

Here, we used a previously established murine model of FGR in order to assess the effect of an adverse *in utero* environment on neonatal and adult immunity. The *Igf-2*^*P0*^ (P_0_) transcript is imprinted and paternally expressed leading to specific loss of placental *Igf2* and growth restriction in the fetuses carrying a P_0_ transcript deletion (16). We then used Drop-Seq to profile the transcriptomes of 7264 cells from neonatal thymi in order to characterize possible immune perturbations at a cellular level.

## Methods

### Mouse tissue preparation

Mice were maintained at Central Biomedical Services in accordance with the UK Home Office, Animals (Scientific Procedures) Act 1986 which mandates ethical review. C57BL/6 dams were mated with Igf-2^P0^ heterozygous males. Mice were genotyped as previously described (35). For neonatal analysis, mice were weighed five days after birth prior to organ collection. For adult mice, body and organ weights were assessed at 8 weeks of age. Spleens were digested with collagenase D (11088866001, Sigma, Gillingham, UK) at 1mg/mL in HBSS (24020091, ThermoFisher, Paisley, UK) for 30 minutes at 37°C and washed with PBS (D8537, Sigma). Single cell thymic suspensions were prepared by maceration of the thymi through a 40 µm strainer. Red blood cell lysis was performed with PharmLyse (555899, BD Biosciences, Wokingham, UK) for 3 minutes at RT for both spleens and thymi. Resultant suspensions were passed through a 40 µm strainer and live cells counted using trypan blue (T8154-20ML, Sigma).

### Flow cytometry

Isolated cells were incubated with antibodies listed in Table S1 on ice for 30 min in PBS prior to washing. Cell types were identified as in Table S2. Samples were processed on a BD LSR Fortessa and the data was analyzed using FlowJo 10. Neonatal and adult samples were analyzed using the same gating strategy (Figures S1 – S3).

### Statistical analysis

For fold change calculations, Igf-2^P0^ mice were compared to litter mate controls with each metric being divided by the mean value obtained from WT litter mates. Outliers were identified within each data-set using the ROUT method in GraphPad Prism where Q = 1. This identified only one outlier (Splenic immune subsets, Figure 1, P_0_) which was subsequently removed. Datasets were tested for normality using the Shapiro-Wilk normality test. Normally distributed data were compared using unpaired two-tailed Student’s t-test while nonparametric data were compared using the Mann-Whitney Rank comparison test. Cell cycle data was analyzed using a two-way ANOVA with Tukey’s multiple comparisons test across cell types or multiple t-tests using Two-stage step-up method of Benjamini, Krieger, and Yekutiele across genotypes.

**Figure 1:**
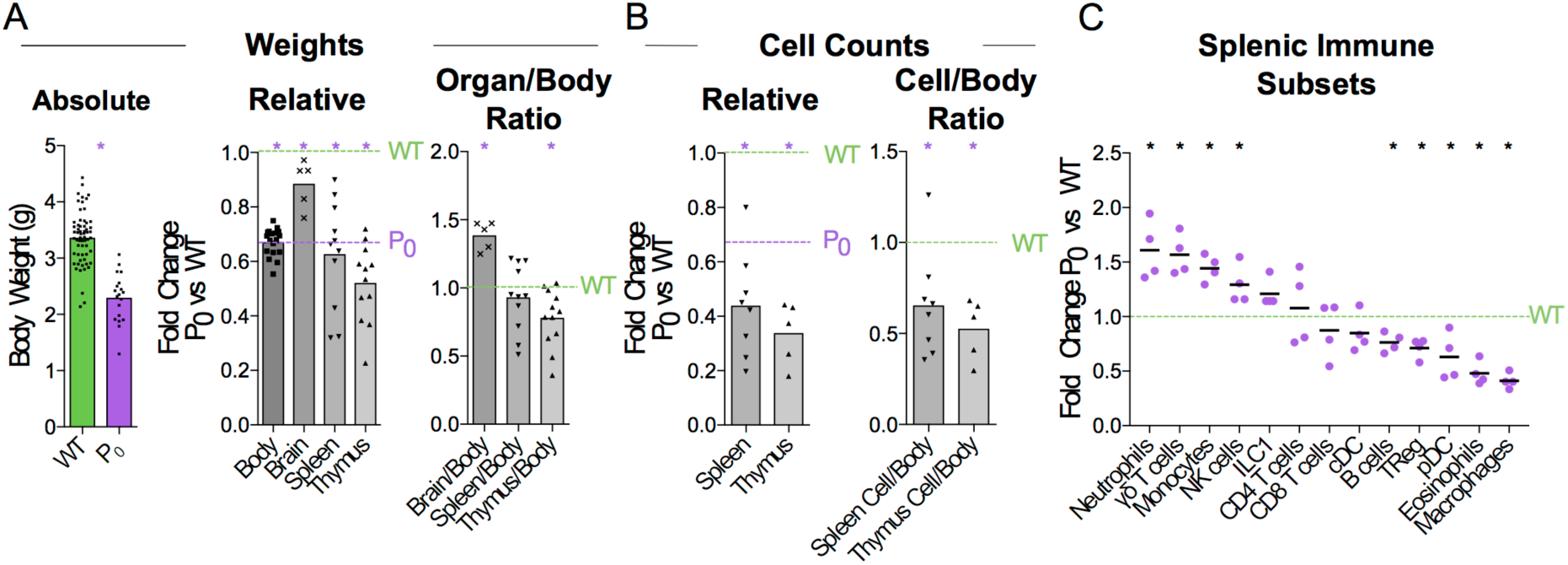
FGR leads to Brain-Sparing at the Expense of Thymus. WT dams were mated with P_0_ males to produce P_0_ and WT offspring. The number of WT and P_0_ animals or organs analyzed for each measurement is given in parentheses. (**A**) Neonates were analyzed five days after birth for absolute body weight (n *=* 61, 19). Relative body and organ weights from P_0_ neonates were compared with littermate controls. Purple dashed line was defined by the average P_0_ body weight, to allow relative comparison of organ weights to both P_0_ and WT. (Relative Body Weight n = 41, 18; Brain n = 7, 5; Spleen n = 32, 11; Thymus n = 28, 12). Organ/Body ratios were compared between P_0_ and WT littermates. (**B**) Absolute cell counts within the spleen and thymus were compared with littermate controls (Spleen Cells n = 16, 8; Thymus Cells n = 7, 5). Cell counts as a ratio to mouse weights were compared with littermate controls. (**C**) Spleen cells were analyzed using flow cytometry and the proportions of the indicated immune cell populations were compared with littermate controls (Spleen Flow Cytometry n = 10, 5, with 1 outlier removed using the ROUT method). Datasets were tested for normality using the Shapiro-Wilk normality test. Normally distributed data were compared using unpaired two-tailed t-tests while nonparametric data were compared using Mann-Whitney Rank comparison tests. For all tests, * indicates P < 0.05.

### Drop-seq

Thymi were dissected from postnatal day (PND) 6 neonates and homogenized in PBS before passing through a 40 µm filter. Cells were counted and diluted to 200,000 cells / mL in PBS/0.1% BSA (A7906-100G, Sigma) for processing according to Drop-Seq protocol (36). Single cells were encapsulated in droplets with primer barcoded beads (MACOSKO-2011-10, ChemGenes, Wilmington, USA). After RNA binding to the oligo-dT portion of the primer, the droplets were broken to pool beads for reverse transcription (EP0743, ThermoFisher). Exonuclease (EN0581, ThermoFisher) treatment removed unused primers from the beads prior to PCR amplification at up to 8,000 beads / tube (∼400 cells/tube). Samples were pooled with up to 4,000 cells / sample and the DNA purified before determining concentration. DNA was fragmented and tagged with Illumina adapter and index sequences using Nextera Sequencing kit (FC-131-109, Illumina, Little Chesterford, UK) and 600 pg DNA/sample. Samples were diluted 1:1 in ddH_2_0 water before double-sided purification with Agencourt Ampure beads (A63881, Beckman Coulter, High Wycombe, UK), using 0.6x beads to remove large fragments and 1x beads twice to remove contaminating primers. Samples were analyzed for size distribution using an Agilent HS DNA kit (5067-4626, Agilent, Stockport UK) and concentration and sequenced on an Illumina HiSeq4000 with an Index read of 8bp, Read 1 of 25 bp and Read 2 of 100 bp. Samples were collected from two separate litters (two WT and one or two P_0_ animals per litter). Single-cell barcoded cDNA was generated for each sample with littermates being processed on the same day. Library preparation was done for all samples on the same day using Nextera indices to distinguish between each neonate. All samples were sequenced together in one lane of an Illumina HiSeq4000 flow cell.

### Bioinformatics analysis

All custom code is freely available on GitHub (https://github.com/darogan/2018_Bacon_Charnock-Jones). Raw Fastq files are demultiplexed (dropseq_demultiplex.sh) using the Nextera indices and then converted to uBAM using PicardTools:FastqToSam (v2.9.0). Quality control, alignment (STAR v020201) gene quantification and final matrix generation were performed using DropSeqTools (v1.12, http://mccarrolllab.com/dropseq). Alignments were performed against the mouse reference genome (mm10 available from http://mccarrolllab.com/dropseq). Initial thresholds of a minimum 200 genes per cell and genes must be present in at least 3 cells were applied. The resulting digital expression matrix (DEM) was imported into Seurat (37) (v2.3.0) for downstream analysis and is split into two scripts. The first, dropseq_seurat_splitDEMs.R, performs the more computationally intensive tasks intended to be run on high performance computers, the Seurat object is saved in Robj format to be imported in to second script, dropseq_seurat_splitDEMs_Plots.R, for differential transcript analysis and figure creation.

Two separate DEMs were calculated for the WT and WT+P_0_ samples. The WT only samples were used to calculate variable genes (FindVariableGenes), which were then used as input to generate the PCA (RunPCA), find clusters (FindClusters) and produce a tSNE (t-distributed stochastic neighbor embedding) visualization (RunTSNE) from the combined WT and P0 sample DEM. FindClusters is run across multiple resolutions (0.2, 0.4, 0.6 0.8 and 1.0), each stored on the Seurat Object. Normalization (NormalizeData), UMI and MT regression (FilterCells) were performed using Seurat including a more stringent threshold of a minimum 300 genes per cell and genes must be present in at least 3 cells was applied. Cell cycle assignments were performed using SCRAN (38) (v1.6.9) on the combined WT+P0 DEM, using an intermediate SingleCellExperiment (v1.0.0) data structure, and then added back to the Seurat Object. Cell cycle genes were regressed out using a subtraction of G2M from S cell cycle scores per cell. The resulting Seurat data object is saved as an RObj for input into the plotting and differential analysis part of the pipeline.

The RObj generated from first dropseq_seurat_splitDEMs.R script are imported into dropseq_seurat_splitDEMs_Plots.R to extract (e.g. with GetCellEmbeddings) the required data for each of the plots in the figures. Custom tSNE plots were generated using ggplot2. Transcript abundance dotplots were generated from AverageExpression extracted from the Seurat object and ggplot2. Cluster trees were generated using clustree (39). Differential transcript analysis was performed by comparing each cluster in turn to all others (FindAllMarkers) and using a log fold change threshold of > 0.7 and adjusted p value < 0.01. The heatmap (pHeatmap), used the same thresholds, and the top 20 genes for each cluster selected.

A custom tool was created to classify whether ribosomal proteins are localized on the surface or are internal to the ribosome structure (https://github.com/darogan/Ribosomal-Protein). The bioinformatics tool scores each ribosomal subunit by its distance from the subunit core. We used this to identify the subunits present on the surface of the ribosome. The tool also provides features to display and colour the subunits in Pymol, and was used to confirm they are accessible on the surface. A further feature links the gene names for the subunits to the nomenclature used in the structure file.

Raw sequencing data have been deposited in the ArrayExpress database at EMBL-EBI under accession number E-MTAB-6945 (https://www.ebi.ac.uk/arrayexpress/experiments/E-MTAB-6945).

## Results

### Placental insufficiency leads to brain sparing at expense of thymus

When crossing wildtype (WT) C57BL/6 females with heterozygous mutant *Igf2* P_0_ males, both WT and P_0_ offspring develop in the WT mother. The P_0_ neonates exhibit poor placental and fetal growth. Animals were assessed five days after birth for body and immune organ weights, with littermates used to determine fold changes (Figure 1A). This time-point was chosen as it is early within the murine neonatal period but provided sufficient tissue. Consistent with previous data(16), the overall body weight was reduced while brain weight was only marginally affected. Spleens of FGR P_0_ neonates were smaller than WT but this change was proportionate to body weight. However, overall cell number was reduced (Figure 1B). Further investigation using flow cytometry revealed a number of perturbed lineages, where neutrophils and macrophages were the most affected by FGR (Figure 1C). The thymus suffered markedly more growth restriction than the spleen with disproportionately reduced size and a nearly 70% reduction in cells compared to littermate controls. Given the overall severity in thymocyte reduction, we used Drop-Seq to investigate the effects of FGR at the single-T-cell level. Within the *Igf-2* locus, neither WT nor P_0_ thymocytes cells contained reads aligned to the P_0_-specific upstream exon U2 (Figure S4), confirming that placental-specific P_0_ was not expressed in T cells. This suggests, therefore, that the observed phenotype was a not result of a direct effect of loss of local *Igf-2* but rather, insufficient placental supply of nutrients. The Igf2 locus contains a number of transcript variants from the placental P_0_ through to P_4 (40)_. We also examined reads mapping to the other isoforms and did not notice any distinct patterns except that in the P_0_ mutant there were a number of reads mapping to an untranscribed region upstream of P_1_.

### Drop-Seq identifies developing thymocytes

To investigate the effect of FGR on T-cell development, we performed Drop-Seq analysis of 4,679 WT and 2,585 P_0_ thymic cells (n = 4 and 3 animals respectively) with an average read depth of 32,399 reads/cell resulting in an average of 994 transcripts per cell (sequencing statistics are in Table S1). We performed a post-hoc power estimate based on our sequencing metrics (41). Our 7,264 cells have a mean reads (per gene per cell) of 5.8 which is well in excess of the minimum threshold of 0.02. This also holds for the WT (6.6) and P_0_ (4.0) experiments if considered separately.

The transcript signatures of each WT cell were used to identify clusters and the genes driving that clustering. We regressed out the signals from cell cycle regulator genes when determining the cluster map, which they otherwise affected due to the intrinsic link between cell cycle and T-cell differentiation (42) (Figure S5). Using the remaining cluster determinants, we mapped both WT and P_0_ cells into 8 clusters of 7,264 single cells (Figure *2*A). Initially, we used known differentiation markers to define these clusters (Figure 2B & C). We identified the known T-cell maturation pathway, beginning with cells containing high levels of *Il2ra* (Double Negative, DN) through to *Cd4/8* containing cells (Double Positive, DP) and ending with high levels of *Ccr7* and *Itm2a* (T_Mature_). In this map dominated by DP cells, we were unfortunately not able to distinguish CD4 and CD8 single positive mature T cells. Bias in identifying clusters was checked using various degrees of resolution, from 0 (identifying only 1 cluster) to 1 (identifying 11) (Figure 2D and Figure S6). This shows distinct DN1 separation even at the lowest resolution but heterogeneous DP subclustering. Regardless of resolution, cells clustered into distinct T-cell populations based upon differentiation status (Figure 2E). Of note, we found that the DP cells were highly heterogeneous and could be divided into multiple clusters, suggesting that the traditional definition using two markers is insufficient to fully characterize these cells. Cells in Cluster 7 were identified as contaminating red blood cells due to high hemoglobin mRNA levels, while Cluster 8 contained macrophages, identified by abundant *Aif1 Lyz2*, Complement and H2 complex transcripts. As both of these clusters contained fewer than 100 cells, we focused our attention on the larger T-cell clusters.

**Figure 2:**
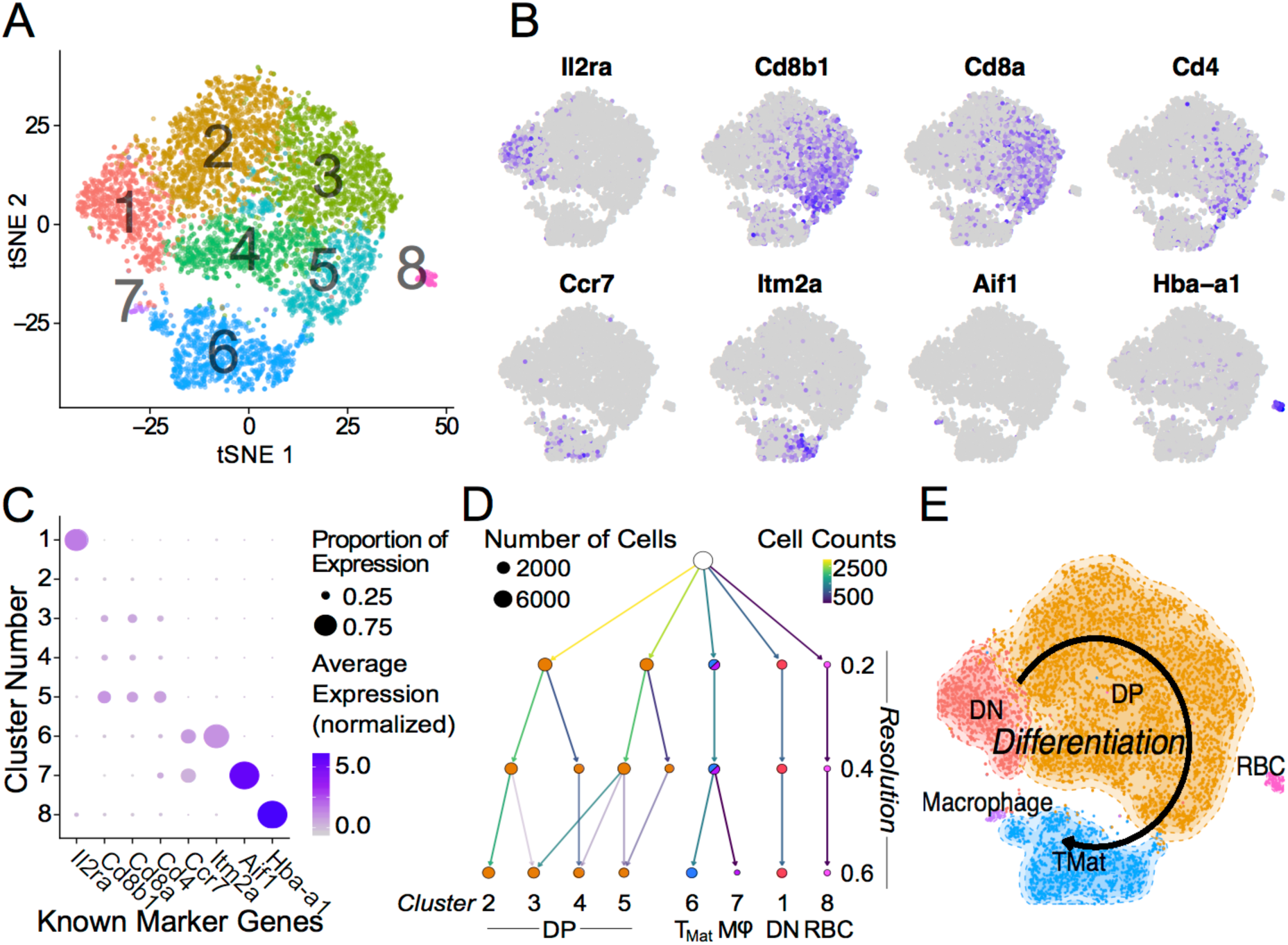
Single Thymic Cell Transcriptomics reveal T-Cell Differentiation Pathway. (**A**) Drop-Seq of 7,264 single-cells from WT and P_0_ thymi were mapped using non-linear dimensional reduction (t-Distributed Stochastic Neighbor Embedding (tSNE)) using WT signatures derived from linear dimensionality reduction (Principal Component Analysis, PCA), with signal from known cell cycle genes regressed. Cells separated into eight clusters, denoted by color and number. Known marker genes were used to identify each cluster. (**B**) Marker transcript levels across cells. Purple indicates detected transcripts. (**C**) Marker transcript levels within each cluster. Average Expression Normalized refers to the amount of transcripts discovered while Percentage Expression denotes the proportion of cells within a given cluster containing the transcript. (**D**) Consistency of sub clustering was assessed at three levels of resolution. (**E**) Overall, these markers yielded a differentiation map from DN in Cluster 1 through to mature T cells in Cluster 6. RBC: Red blood cell.

### Drop-Seq identifies transcript signatures in T-cell maturation

To further characterize these cell clusters, we identified the top marker genes in each cluster by performing a differential analysis of each cluster compared to all others in an unsupervised manner (adjusted P value <0.01; Table S2). These genes included both known T cell specific markers such as Cd8 and Ccr9 and cytoskeletal components (Figure 3). We prevented cell cycle genes driving the clustering (by regression), but in the differential expression analysis the unregressed data were used. A number of cell cycle regulators were found to differ between mapped clusters, demonstrating a clear connection between cellular phenotype and the stage of the cell cycle. Cluster 1, containing the DN cells, was characterized by early T-cell markers *Il2ra* and *Hes1*. *Tcr*γ*-C1* and *C2* were also found within these cells, likely because only DN cells contain precursor Tγδ cells. Cluster 2, the early DP cluster, contained T-cell genes *Granzyme A* and *Ctage5*, as well as *Ranbp1 Nolc1*, and *Hspd1*, which are associated with nuclear pore transport, rRNA processing and protein folding, respectively. *Plac8* was also found, which was previously implicated in T-cell development (43) but here specifically found within DN cells. Clusters 3 to 5, which we have identified as increasingly mature DP cells, contained high levels of *Tarpp* (also known as *Arpp21*), *Rag1* and *Rag2*, whose proteins are present during TCR rearrangements. During T-cell positive selection, *Tarpp* levels are reduced (44), which is shown here as the T_Mat_ cluster contains few Tarpp transcripts. These cells also showed increasing levels of *Ccr9*, the CCL25 receptor expressed on late DN and DP T cells. These known T-cell markers were found alongside *Btg2*, a cell cycle inhibitor, and *Rmnd5a*, which is necessary for mitotic division. Clusters 2 & 3 both contained *Rrm2 Tyms*, and *Esco2*, which synthesize nucleotides and bind sister chromatids, respectively, and are required for S phase of the cell cycle. The variations of Clusters 2 through 5 then are likely in part due to progressive rounds of proliferation as the T cells mature.

**Figure 3:**
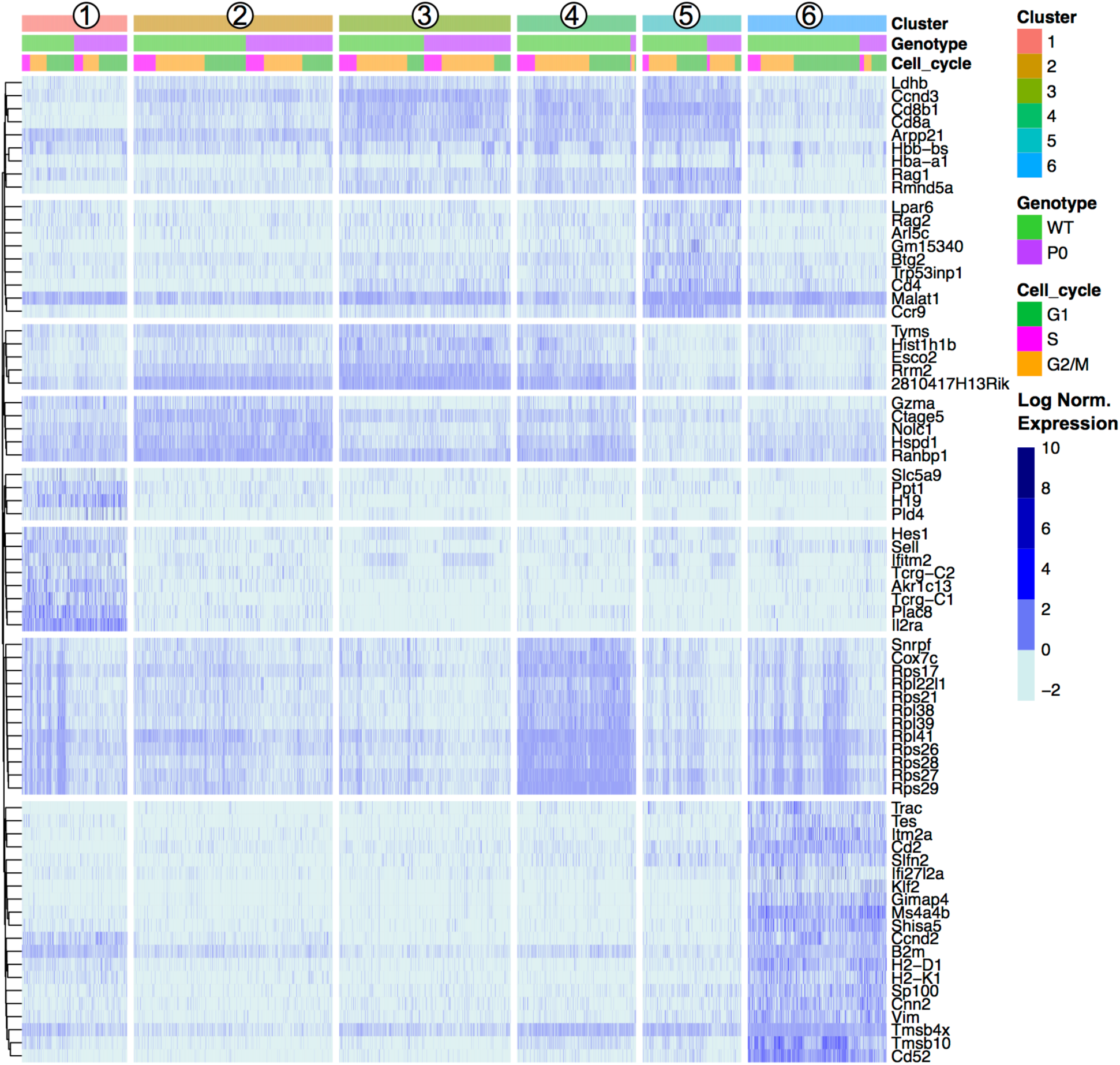
Unsupervised Clustering Driven by T-Cell, Cell Cycle, and Ribosomal Genes. Transcript levels of each cluster were compared with the remaining overall cell population. The top 20 genes with an absolute fold change above 5 and an adjusted p value < 0.01 for each cluster were visualized in a heat map and ordered by genotype (WT vs P_0_) followed by cell cycle score (G1, G2/M, or S). The top 20 genes for each of the six clusters results 120 genes, reduced to 72 unique entries. Each column represents one cell. Signal intensities are shown on a log_2_ scale. The complete gene lists are provided in Dataset S1.

The mature lymphocyte markers *Itm2a* and *Cd52* are specific to Cluster 6, alongside anumber of cytoskeletal (*Cd2 Tmsb4x*, and *Tmsb10*). Negative growth regulators *Slfn2 Gimap4 Ms4a4b*, and *Shisa5* are also present in high levels in the T_Mat_ cluster, likely involved with positive and/or negative selection. Interestingly, the DN Cluster 1, DP Cluster 4, and T_Mat_ Cluster 6 contained heterogeneous mixtures of cells with high and low levels of transcripts coding for ribosomal proteins.

### Cell cycle and ribosomal protein structure drive thymocyte maturation

We further investigated the trends in cell cycle and ribosomal protein genes. Although we prevented known cell cycle genes from determining the cluster map, we did not remove these genes from the transcript profile of each cell. When comparing transcript signatures between clusters, we detected differences in cell cycle stage, likely due to a link between cell type and proliferation state (See Figure S5 for cluster mapping with and without cell cycle regression). We scored each cell for its G_1_, S, or G_2_/M phase transcript signature (Figure 4A) and compared cycle phase between cell types (Figure 4B). DN and T_Mat_ stages had lower levels of proliferation than the DP stage. There were no differences in cell cycle overall between WT and P_0_ genotypes (Figure 4C).

**Figure 4:**
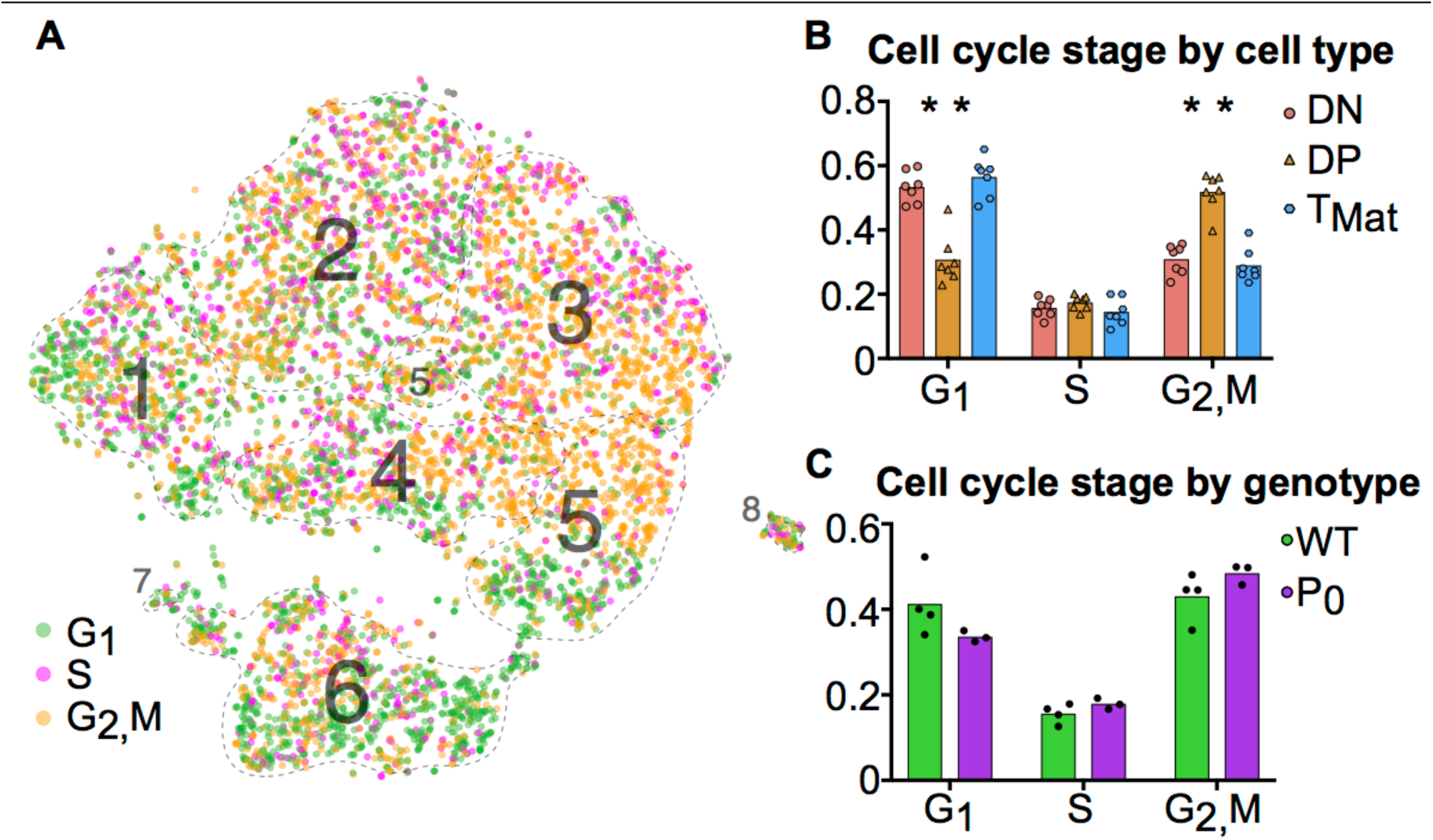
T-cell maturation and cell cycle stage are linked. (**A**) Individual cells were scored for their cell cycle genes (G_1_, S and G_2_/M, as described in Methods). (**B**) The proportions of these cells within each cell population were compared using student t-tests (DN, Cluster 1; DP, Clusters 2-5; T_Mat_, Cluster 6). (**C**) The proportion of WT and P_0_ cells in each phase were compared. (* indicates P < 0.05; n = 7).

A differential transcript analysis for each cluster against all others revealed that within the DP stage, ribosomal protein transcripts were varied, most notably between Clusters 3 and 4. We made the same observation of a differential ribosomal protein transcript population in the unsupervised clustering in Figure 3. The ribosomal protein mRNAs that significantly changed in Cluster 3 were compared against those found in Cluster 4 (Figure 5A). In each case, cells in Cluster 3 were notable for low ribosomal protein transcript abundance while in Cluster 4 such transcripts were increased. The majority of the transcripts (9 out of 13, 69%) found by this comparison between clusters coded for proteins that contributed to the ribosome surface. Surface proteins are more likely to play a functional or regulatory role than internal proteins. The protein subunits of a ribosome are known to play a key role in directing translation of specific mRNA pools (45). Looking at the overall cluster mapping, we found that the ribosomal transcripts appeared in waves dominating Cluster 4 and dividing Cluster 6 (Figure 5B,C, Movie S1).

**Figure 5:**
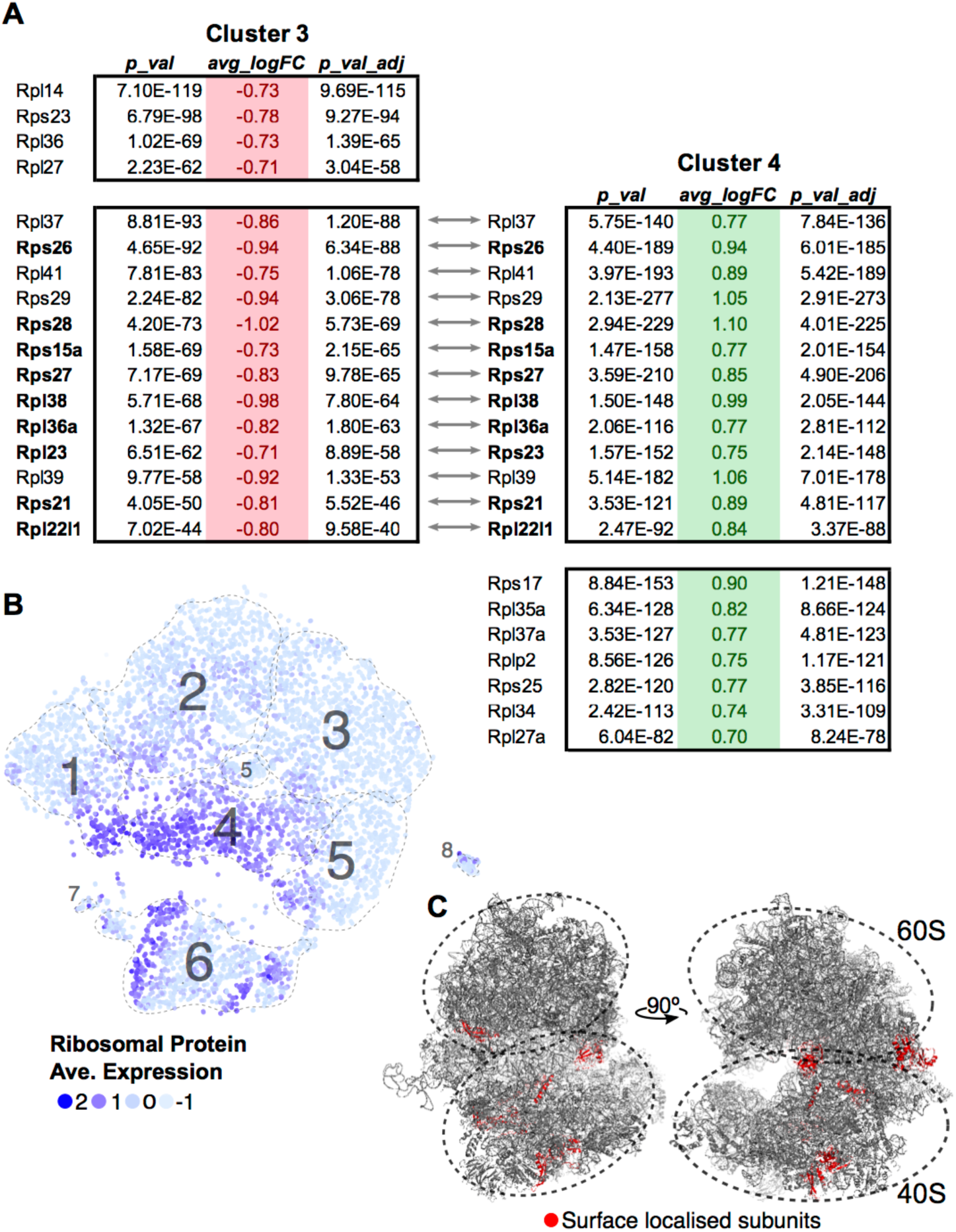
Ribosomal protein transcripts distinguish T-cell maturation steps. (**A**) Significant changes (p < 0.01; log_10_ Fold Change > 0.7) in ribosomal protein transcripts within Cluster 3 and Cluster 4 were compared. Ribosomal genes in bold code for proteins found on the surface of the ribosome. (**B**) Ribosomal protein transcript levels are displayed across cells using average expression values for the 13 genes identified by comparing Clusters 3 and 4. (C) Ribosomal genes identified to be localized on the surface of the ribosome are mapped onto the ribosome structure (red, PDB:6EK0). Supplemental Movie 1 visualises the surface ribosome genes identified on the structure as it rotates on the y-axis.

### FGR prevents thymocyte maturation

Having identified each cluster and compared the top gene signatures, we then compared the WT and P_0_ thymocytes within each cluster (Figure 6A). Even after accounting for the higher number of WT cells present, we still found a higher proportion of DN cells in the P_0_ thymus than WT (Figures 6B & C). The opposite held true with the late stage T_Mat_ cluster, as there were proportionally more T_Mat_ cells in the WT thymus than the P_0_. Overall, there was a general skew of P_0_ cells towards earlier, more immature clusters in the DP pathway, while the WT thymus favored the later stage clusters. Using flow cytometry for surface markers, we detected discrete increases in DN cells at the expense of mature CD4+ cells in the P_0_ (Figure 6D). Within the smaller DN cell population, there were proportionally more DN3 cells in the P_0_ over the WT, which is noteworthy given that rearrangement of V and D chains as well as β selection occur at this stage in T-cell differentiation.

**Figure 6:**
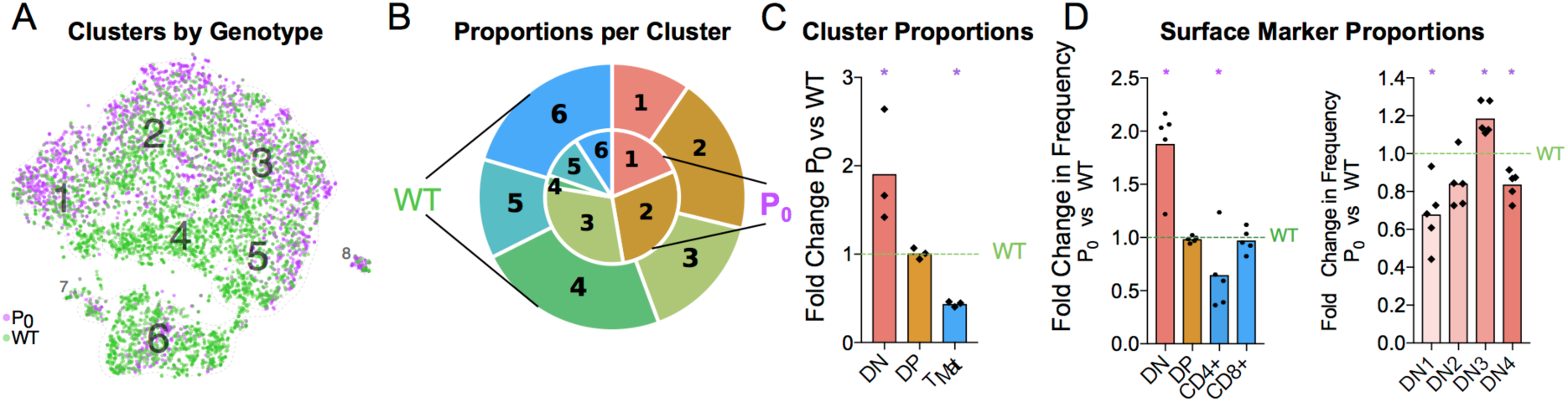
FGR Leads to T-Cell Differentiation Delay. (**A**) Single-cell map colored by genotype with previous cluster numbering. Red: P_0_, 2,585 cells, *n* = 3. Teal: WT, 4,679 cells, (*n* = 4). (**B**) Visual representation of the proportions of cells in each genotype per cluster, averaged from each trial and labelled for each cluster. Clusters are labelled by number, from the DN T-cell stage in Cluster 1 to T_Mat_ in Cluster 7. P_0_ is in the inner circle, while WT is in the outer circle. (**C**) Proportions of cells within each cell population as determined by Drop-Seq were compared to WT levels. (**D**) Cells were classified by their surface markers using flow cytometry as DN, DP, CD4+ or CD8+ cells and proportions were compared with littermate controls (*n* = 9, 5). (**E**) Within the DN population, cells were classified from DN1 to DN4 based on their CD25 and CD44 status. These proportions were then compared with littermate controls. Comparisons were calculated using Student’s t test with the exception of the Cluster 5 (**C**), DN and DN3 (**D**) subsets, which failed to pass the Shapiro-Wilk test for normality and were instead analyzed using Mann-Whitney Rank comparison tests (* indicates P < 0.05).

### FGR Transcript Signatures

Given the apparent waves of ribosomal transcripts appearing in Figure 3, we looked more deeply within clusters to determine whether FGR changed transcript signatures of seemingly equivalent cell types (Figure 7, Supplemental Tables 7A-E). We first identified genes that distinguished major cell type (DN vs DP or DP vs T_Mat_, Figure 7A & B, individual cluster comparisons in Figure S7, Supplemental Tables 7A1-15). As expected, *Cd8* and *Cd4* genes dominated for the DP to DN comparison, whilst Cd52 and Itm2a distinguished T_Mat_ from DP in this unsupervised comparison. We then looked within clusters for changes due to FGR (Figures 7C-E, individual cluster comparisons in Figure S8, Supplemental Tables 8A1-6). Within each cluster, ribosomal protein transcripts were found at significantly higher levels in the WT over the P_0_. *Xist* was also found, which indicates the presence of a female in the P_0_ group (Figure S9 shows the consistent overlap of the female P_0_ with the male P_0_ neonatal T-cells indicating that sex has little effect on these cells). Interestingly, *Igf2* was present in lower levels in the WT DN and T_Mat_ clusters, note that this refers to total *Igf2* rather than the placental specific isoform. We then investigated overarching trends across clusters by comparing genes which differ between WT and P_0_ cells, within each cluster to another cluster (Figures 7F & G). The ribosomal protein transcripts aligned clearly, demonstrating an overall trend as opposed to a cluster-specific phenomenon.

**Figure 7:**
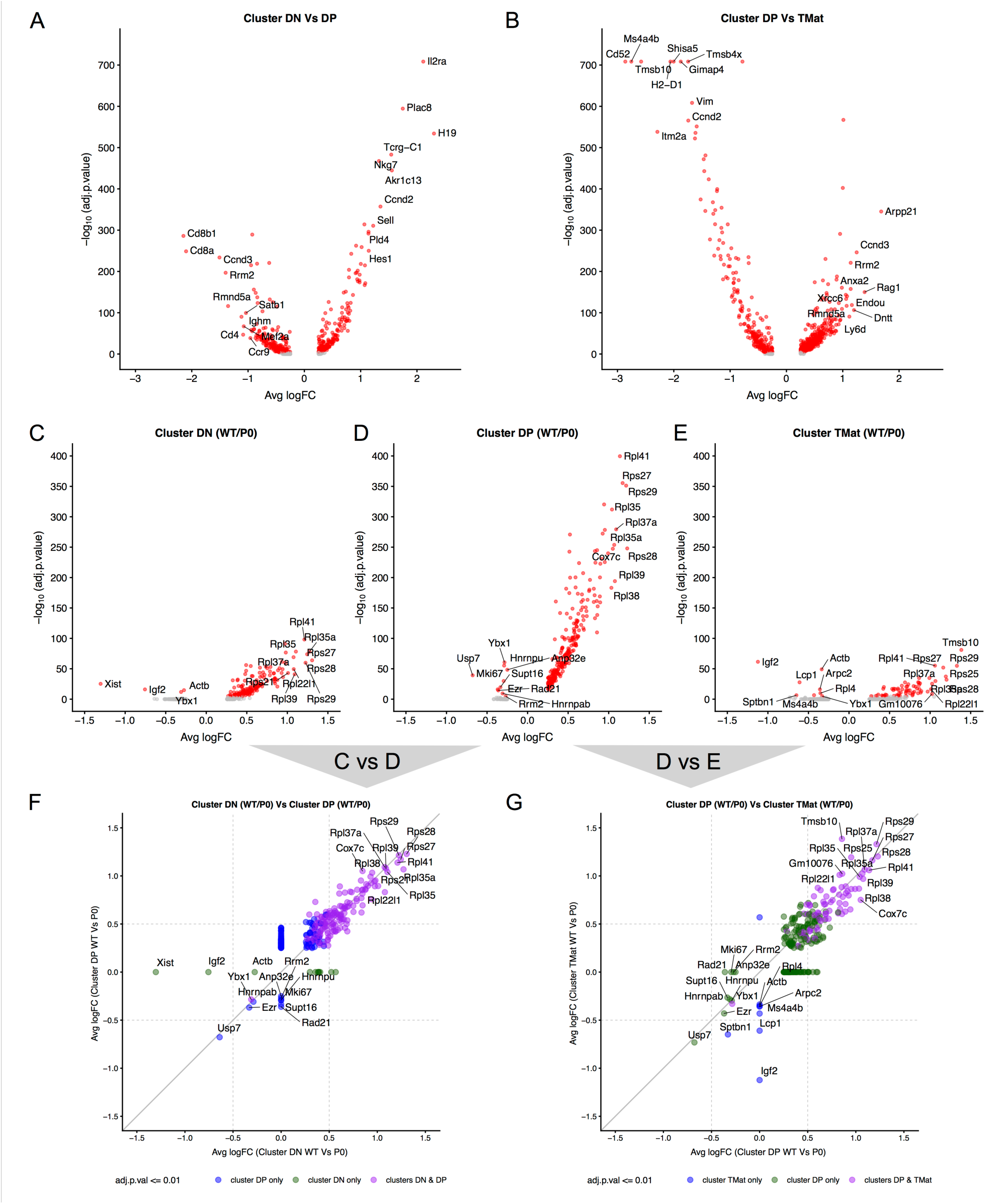
Differential transcript levels between identified cell type clusters and within cluster WT/P_0_ differences. (**A, B**) Volcano plots for differential transcript levels between the DN / DP and DP / T_Mat_ cell type clusters. The top 10 genes are labelled for +/- average log fold change. (**C, D, E**) Within cluster, WT vs P_0_ cell transcript level comparisons revealing the ribosomal protein genes as the dominant set of genes especially evident within the DP cell type. (**F,G**) Comparison of the transcript level fold change between DN / DP and DP / T_Mat_ cell type clusters shows a similar enrichment for the ribosomal protein genes (purple).

### Immune defect lasts into adulthood

Thymus size is correlated with thymus function (46). Humans with DiGeorge syndrome have little or no thymus, resulting in a significant immunodeficiency (47,48). We wondered whether the maturation delay in the thymus of the 6-day old neonate would affect the immune cell population in the adult. Infants with FGR exhibit increased growth and ‘catch up’ relative to their unrestricted counterparts (49). Adult male mice displayed signs of catch-up growth, although they were still found to be significantly lighter than their littermate controls (Figure 8A). However, the spleen and thymus weights were not different from littermate controls. Despite this, the cellularity of the thymus was still significantly reduced (58% ±.19%, P = 0.019, Figure 8B). However, as both DN and CD4+ lineages recovered to WT proportions the long-term effects of poor placentation on cell distributions in the adult thymus are less clear (Figure 8C). While we did not evaluate the repertoire diversity or clonality, we did observe that DN subtypes were statistically distinct from WT (Figure 8D).

**Figure 8:**
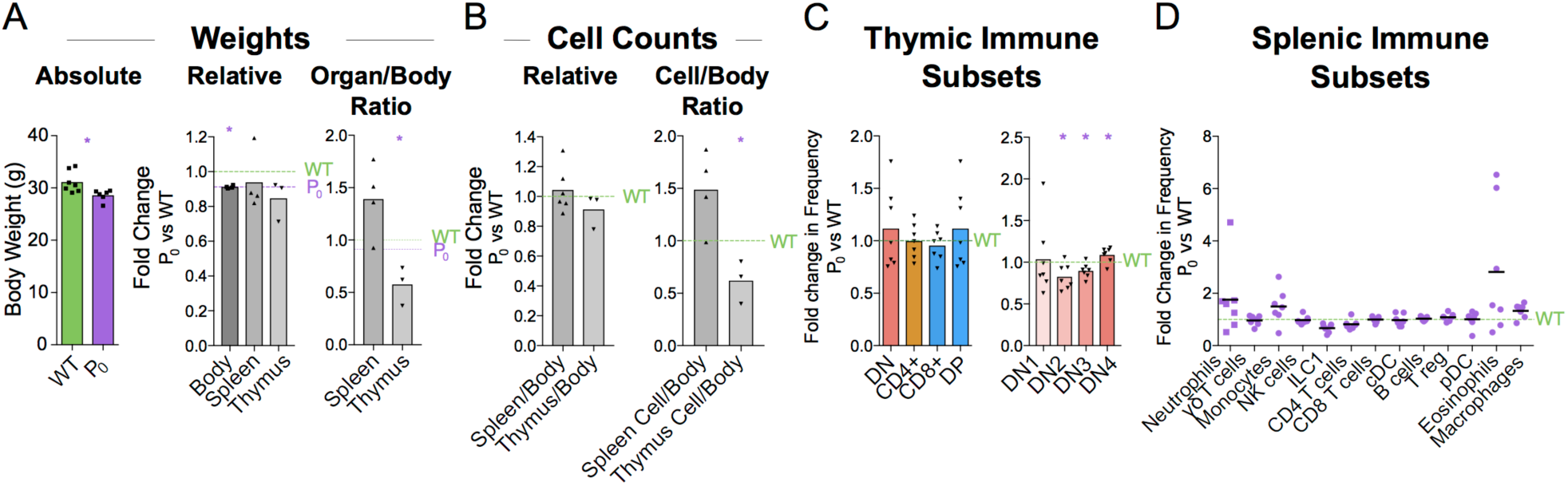
Immune cell deficiency persists into adulthood. (**A**) Absolute weights of 8 week male P_0_ and WT were compared (n = 7, 6). Body (n = 7, 4), spleen (n = 7, 4), and thymus (n = 5, 3) weights were compared with littermate controls. Spleen/body and thymus/body weight ratios were also compared relative to littermate controls to account for lower overall body weight in the P_0_ mice. (**B**) Total cells within the spleen (*n* = 4, 4) and thymus (n = 5,3) were compared directly with littermate controls as well as relative to overall mouse weight. (**C**) Surface markers were used to classify T cells within the thymus and the proportions were compared to littermate controls (n = 8, 7). (**D**) Splenic immune subset proportions were determined from flow cytometry and compared with littermate controls (n = 5, 7). Datasets were tested for normality using the Shapiro-Wilk normality test. Normal data were compared using unpaired two-tailed t-tests while nonparametric data were compared using Mann-Whitney Rank comparison tests. For all tests, * indicates P < 0.05.

## Discussion

We present the first single-cell analysis of the neonatal thymus, in the context of a disease model for FGR. Recently, interest in fetal immunity has led to the generation of a cell atlas for the developing thymus (50). Here, we add to this atlas with an insight into thymic development after birth. We found a number of genes in common with these early developmental snapshots, such as *H19* and *Plac8,* but were further able to resolve multiple developing T-cell populations due to the seemingly increased thymic activity after birth.

The “fetal programming of adult disease” theory has largely been associated with noncommunicable diseases such as obesity or heart disease. Within this wider context, we present the first genetic model suggesting that an adverse *in utero* environment perturbs immune cell development. This may lead to poor immune function later in life and consequently a higher risk of communicable diseases.

Our study has identified a novel sequela of FGR in the mouse: a lasting deficit in T cells, resulting from a maturation delay found in the growth restricted neonate. At the cellular level, there were more DN cells and a skew towards early DP cells. The high heterogeneity of DP cells found here suggests that current definitions of DP are not specific enough given the large proportion of the thymic T cells in this category. Our 26 supplemental tables from the single-cell dataset provide numerous candidates for future investigation of subcluster markers and some of these may be functionally relevant to the differentiation process. Of particular note is the high number of ribosomal protein transcripts driving clustering within this group. Ribosome biogenesis has recently been shown to play a key role in stem cell differentiation (51) and translation of specific pools of mRNAs depends on the ribosomal protein composition (45). This is particularly noticeable with defects in specific ribosomal protein genes, which can lead to discrete effects such as an erythropoiesis block in hematopoietic stem cells (52) or a loss of body hair (53), as opposed to an expected generalized protein deficit. Clusters 1, 4 & 6 - comprised of DN, late DP and maturing T cells - seem to subcluster based on ribosomal protein transcript levels. The ribosomal proteins encoded by these transcripts, located largely on the outside of the ribosome, are greatly reduced in the P_0_ mice. It is interesting to speculate that within this T-cell differentiation pathway, ribosome biogenesis, and hence function, plays a role in directing maturation.

The clusters were defined by transcript signatures after excluding cell cycle regulators. Nevertheless, distinct cell cycle signatures were present within the clusters, confirming that proliferation and T-cell maturation are inexorably linked, which is consistent with a similar finding in specifically CD4+ T-cell differentiation (42).

How this early phenotype leads to low thymic cellularity in the adult remains unclear. Infants with FGR typically undergo ‘catch-up growth’ to reach normal or near normal weight. This was the case here for all parameters except thymus cellularity. T-cell maturation, is linked at a molecular level to cell cycling because RAG2 is degraded during G2, S, and M phases (54). Perhaps thymic ‘catch-up growth’ and T cell diversification and maturation are incompatible because proliferation inhibits T-cell maturation. Indeed, the similar cell-cycle profiles of the P_0_ and WT thymic cells is indicative of a lack of catch-up growth in the P_0_ thymus. It is also possible that catch-up growth cannot occur because the thymus is programmed to involute after birth, rather than expand. In either case, growth restriction early in life would be irreversible: the number of T-cell clones generated in the neonatal thymus restricts the number of T-cell clones present in the adult.

In both mice and humans, thymus output declines with age, with the memory T cells dominating the peripheral pool in old age (55). The diversity and clonality of naive T cell also change with age (56). Immune function declines with age and there is an increased risk of infection and malignancy late in life (46). Genetically manipulated mice with enlarged thymi have increased thymic activity and enhanced recovery from γ irradiation (57). In contrast, patients with DiGeorge syndromes where the thymus is small display immune vulnerability early in life (47,48). Thus, a thymic deficit early in life could have dramatic effects on adult immunity which would only become exacerbated by normal thymic involution.Overall, we have identified key immune phenotypes caused by placental insufficiency. Future studies will ultimately determine the role of FGR in both neonatal and adult immunity. Nevertheless, we have shown a distinct and negative impact of FGR on immune cells in the mouse.

## Acknowledgements

We thank Dr. Miguel Constância for provision of the Igf-2^P0^ mouse line.

## Author contributions statement

FC, JK, DC, and WB contributed conception and design of the study; WB, JK, DH and AK performed the research. RH and WB analyzed the data. RH contributed novel bioinformatic tools. ZY and CA contributed microfluidic equipment and expertise. WB wrote the first draft of the manuscript; RH and DC wrote sections of the manuscript all authors contributed to manuscript revision, read and approved the submitted version.

## Data Availability Statement

The single-cell datasets generated for this study can be found in the ArrayExpress database at EMBL-EBI under accession number E-MTAB-6945, https://www.ebi.ac.uk/arrayexpress/experiments/E-MTAB-6945. All other raw data supporting the conclusions of this manuscript will be made available by the authors, without undue reservation, to any qualified researcher.

## Supplementary Material

### Dataset S1

Transcript markers distinguishing each cluster using an adjusted p value < 0.01 are shown here.

T-Cell.SupplementalFiles.zip

Files contained in the zip file. File names correspond to their associated figure in the manuscript

7A - T-Cell.Table.DN.vs.DP.adjp.0.01.csv

7B - T-Cell.Table.DP.vs.TMat.adjp.0.01.csv

7C - T-Cell.Table.WT_DN.vs.P0_DN.adjp.0.01.csv

7D - T-Cell.Table.WT_DP.vs.P0_DP.adjp.0.01.csv

7E - T-Cell.Table.WT_TMat.vs.P0_TMat.adjp.0.01.csv

S7A.01 - T-Cell.Table.1.vs.2.adjp.0.01.csv

S7A.02 - T-Cell.Table.1.vs.3.adjp.0.01.csv

S7A.03 - T-Cell.Table.1.vs.4.adjp.0.01.csv

S7A.04 - T-Cell.Table.1.vs.5.adjp.0.01.csv

S7A.05 - T-Cell.Table.1.vs.6.adjp.0.01.csv

S7A.06 - T-Cell.Table.2.vs.3.adjp.0.01.csv

S7A.07 - T-Cell.Table.2.vs.4.adjp.0.01.csv

S7A.08 - T-Cell.Table.2.vs.5.adjp.0.01.csv

S7A.09 - T-Cell.Table.2.vs.6.adjp.0.01.csv

S7A.10 - T-Cell.Table.3.vs.4.adjp.0.01.csv

S7A.11 - T-Cell.Table.3.vs.5.adjp.0.01.csv

S7A.12 - T-Cell.Table.3.vs.6.adjp.0.01.csv

S7A.13 - T-Cell.Table.4.vs.5.adjp.0.01.csv

S7A.14 - T-Cell.Table.4.vs.6.adjp.0.01.csv

S7A.15 - T-Cell.Table.5.vs.6.adjp.0.01.csv

S8A.1 - T-Cell.Table.WT_1.vs.P0_1.adjp.0.01.csv

S8A.2 - T-Cell.Table.WT_2.vs.P0_2.adjp.0.01.csv

S8A.3 - T-Cell.Table.WT_3.vs.P0_3.adjp.0.01.csv

S8A.4 - T-Cell.Table.WT_4.vs.P0_4.adjp.0.01.csv

S8A.5 - T-Cell.Table.WT_5.vs.P0_5.adjp.0.01.csv

S8A.6 - T-Cell.Table.WT_6.vs.P0_6.adjp.0.01.csv

### Movie S1

5 - T-Cell.RibosomeMovie.mpg

**Movie S1**: Ribosomal genes identified to be localized on the surface of the ribosome are mapped onto the ribosome structure (red, PDB:6EK0) with a rotation on the y-axis.

---

**Figure S1.**
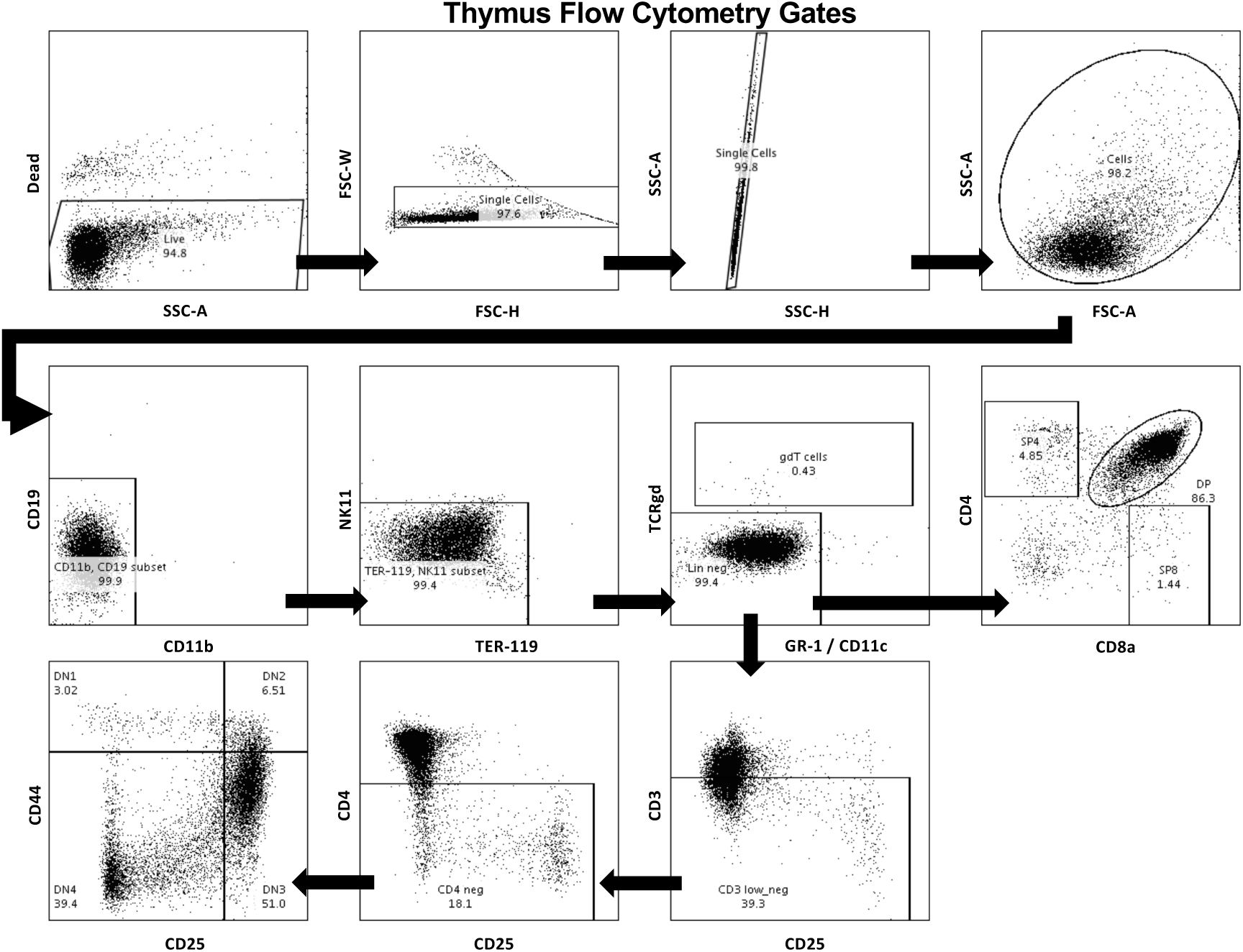
Representative flow plots from a neonatal thymus. The first row demonstrates how we gated for living, single cells, while the remaining rows show our gating for DN, DP, CD4+ and CD8+ T-cells as listed in detail in Table S2. Samples were processed on a BD LSR Fortessa and the data was analyzed using FlowJo 10.

**Figure S2.**
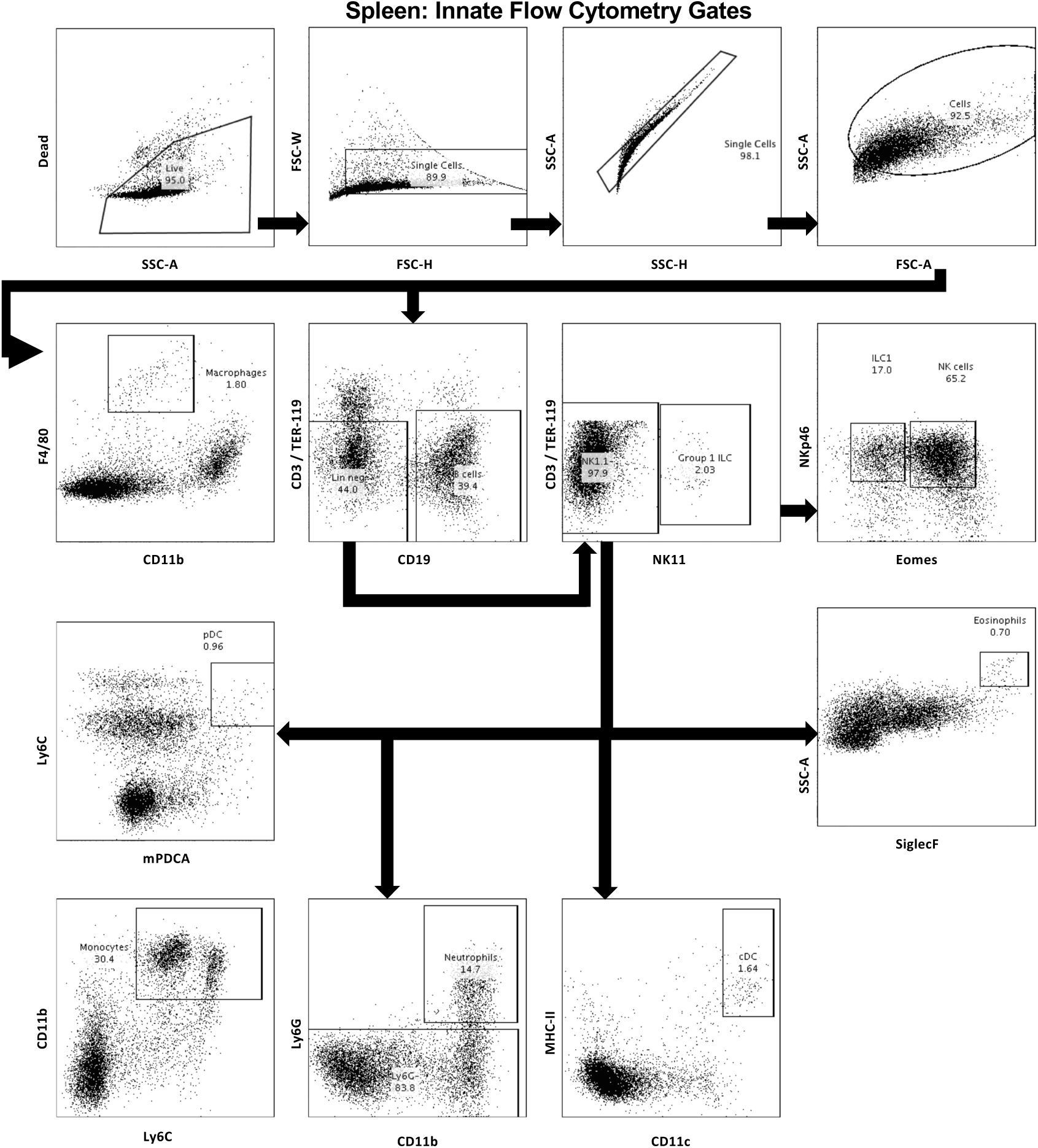
Representative flow plots for innate immune cells from a neonatal spleen. The first row demonstrates how we gated for living, single cells, while the remaining rows show our gating for B-cells and the wide variety of innate immune cells as listed in detail in Table S2. Samples were processed on a BD LSR Fortessa and the data was analyzed using FlowJo 10.

**Figure S3.**
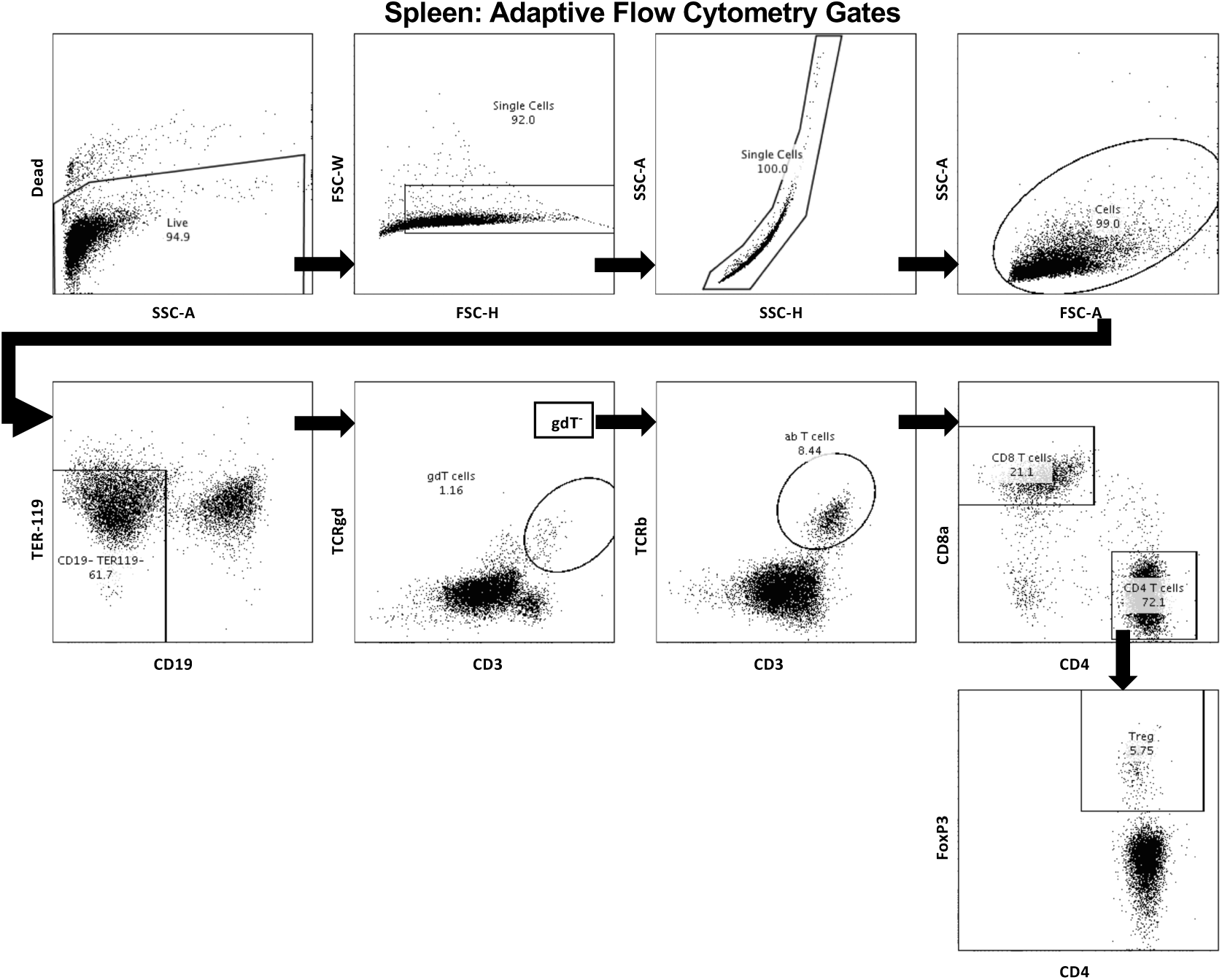
Representative flow plots for adaptive immune cells from a neonatal spleen. The first row demonstrates how we gated for living, single cells, while the remaining rows show our gating for various T-cell types as listed in detail in Table S2. Samples were processed on a BD LSR Fortessa and the data was analyzed using FlowJo 10.

**Figure S4.**
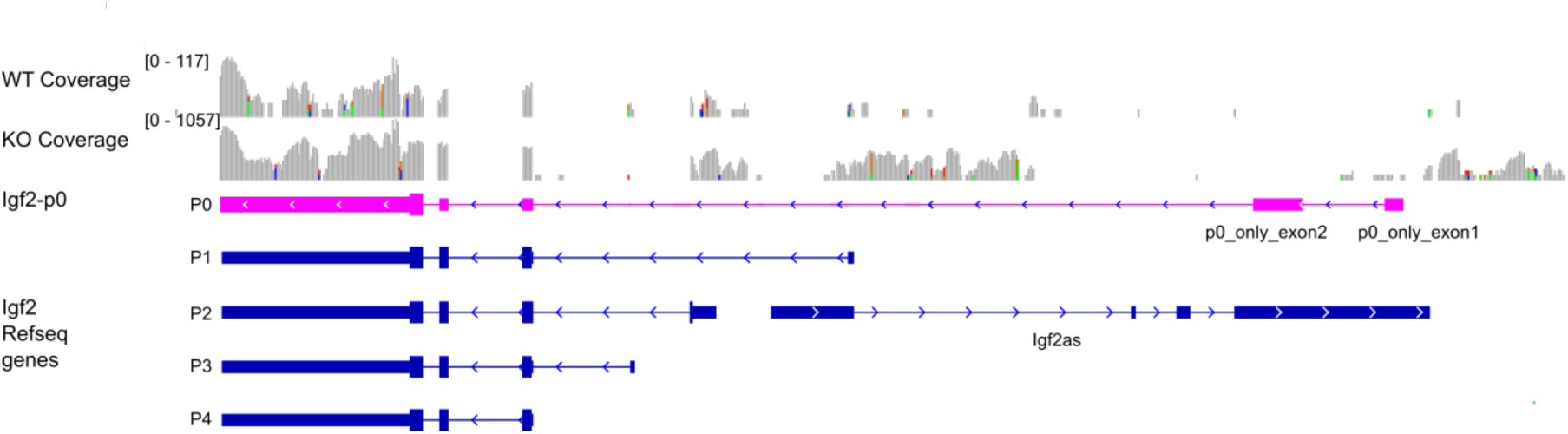
P_0_ *Igf-2* transcript not found in thymus. Total reads from both WT and P0 single-cell transcript sequencing were visualized within the *Igf-2* locus. Transcript isoforms from P0 to P4 are shown. All isoforms yield the same translated product.

**Figure S5.**
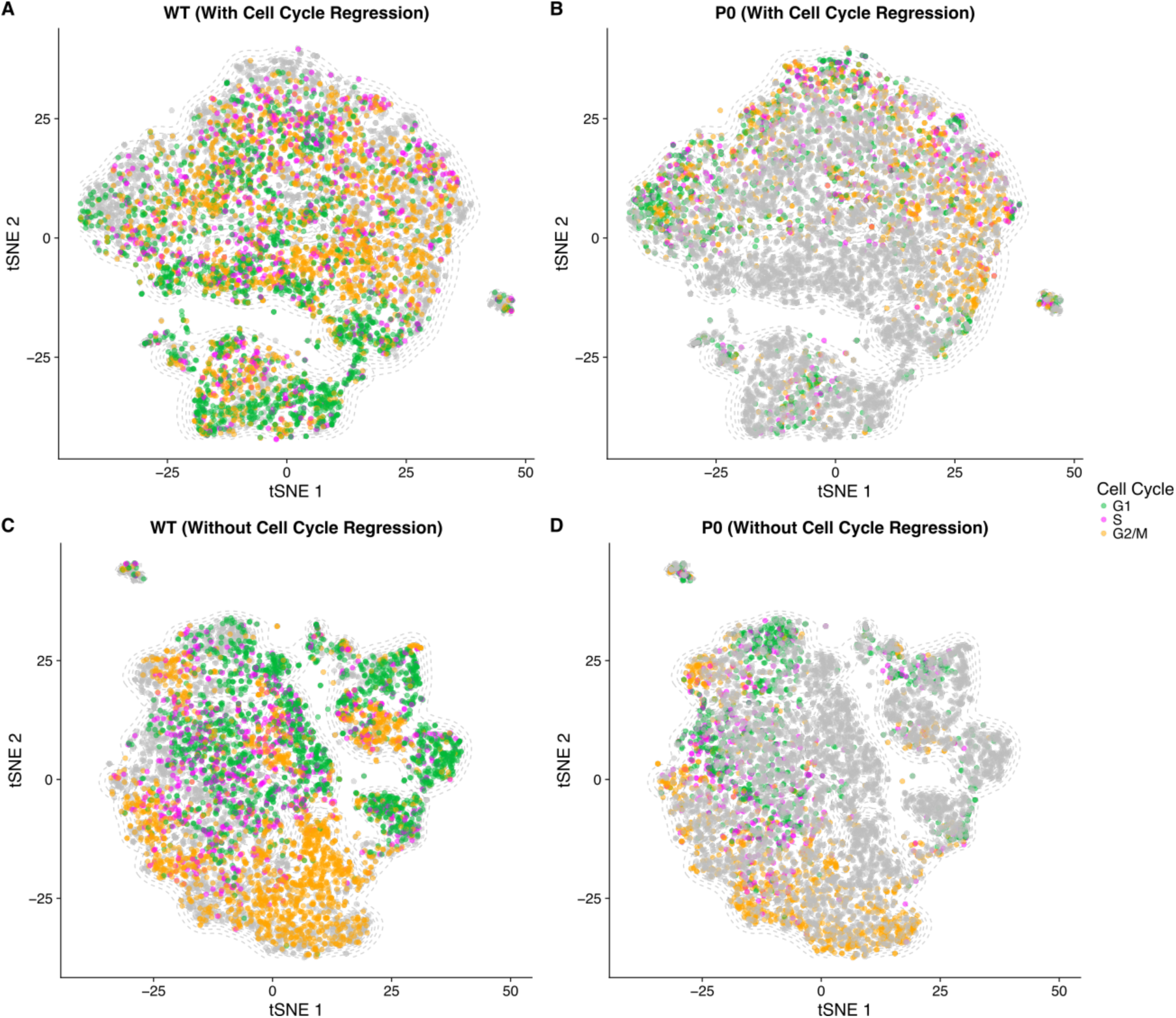
Effect of Cell Cycle Regression on Cluster Mapping. Cells were clustered before and after regressing cell cycle determinants. (**A**) WT and (**B**) P0 cells with cell cycle regression, as compared with (**C**) WT and (**D**) P0 cells mapped without cell cycle regression. Cells are colored by genotype (grey if the opposing genotype) and cell cycle phase (G1: Green, S: Pink; G2/M: Orange).

**Figure S6.**
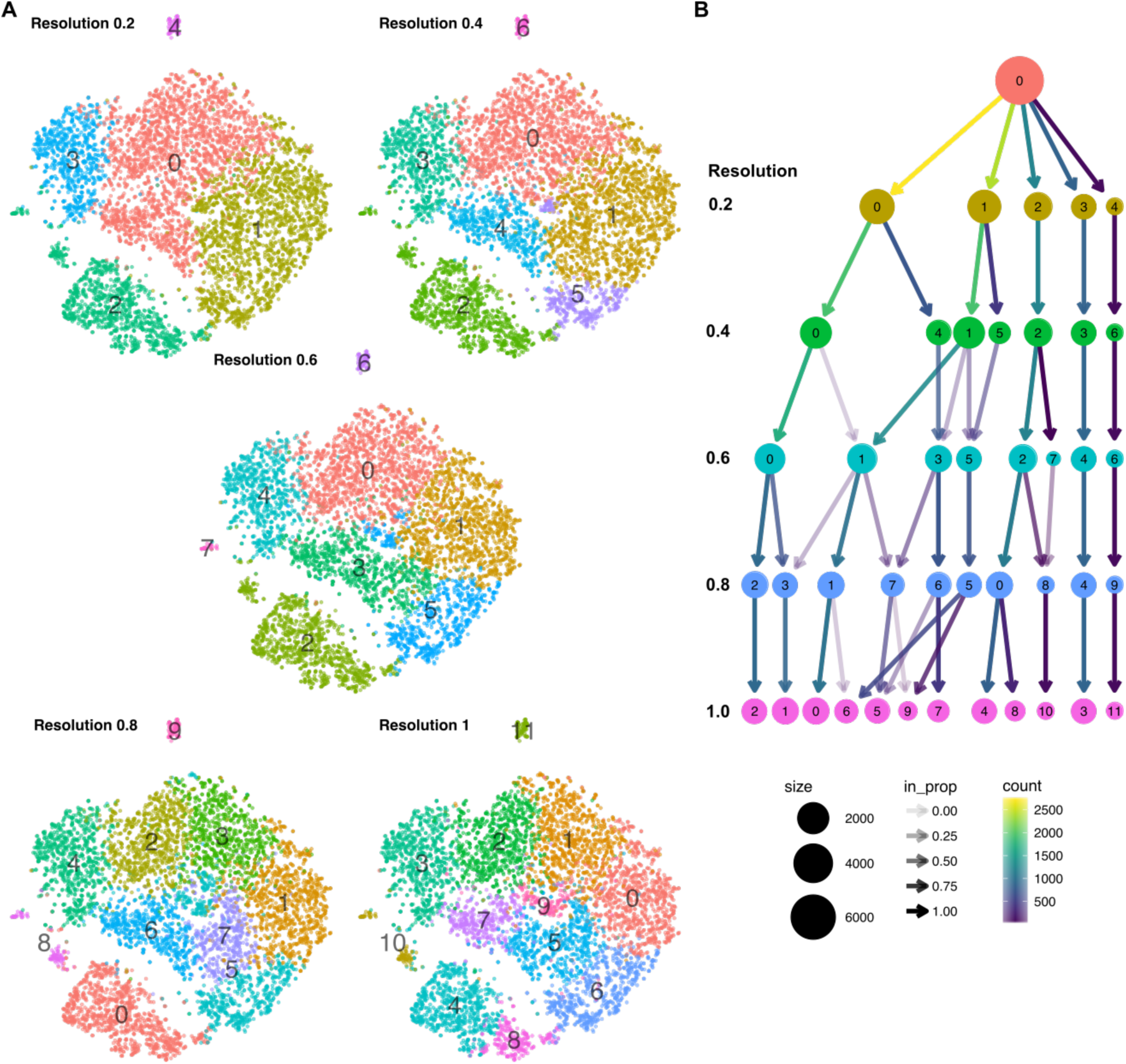
Resolution Effects on Cluster Calling. (**A**) In relation to Figure 2, an increase in resolution of cluster calling leads to increased numbers of clusters. (**B**) The cluster tree diagram shows how consistently clusters are called, for instance the clusters at the right (first row, numbers 3 & 4) are unique clusters no matter what resolution is used. At higher resolutions clusters divide and cells from one or more of the lower resolution clusters contribute.

**Figure S7:**
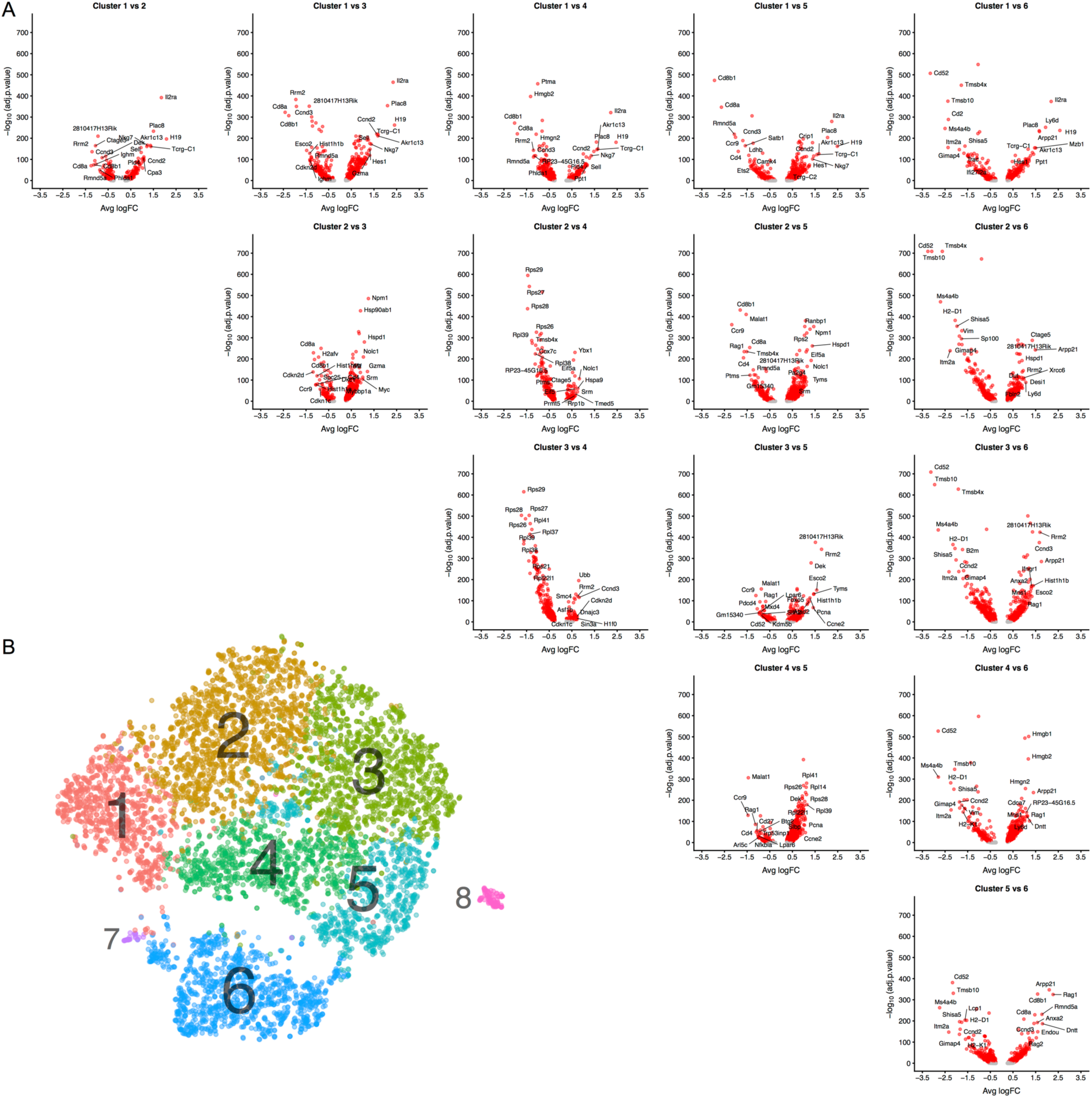
Differential transcript levels between the six largest clusters (1-6). (**A**) Volcano plots for differential transcript levels between each pair of clusters. The identified cluster marker genes were used to assign cell types. The top 10 genes are labelled for +/- average log fold change. (**B**) The clusters compared as shown on the tSNE.

**Figure S8:**
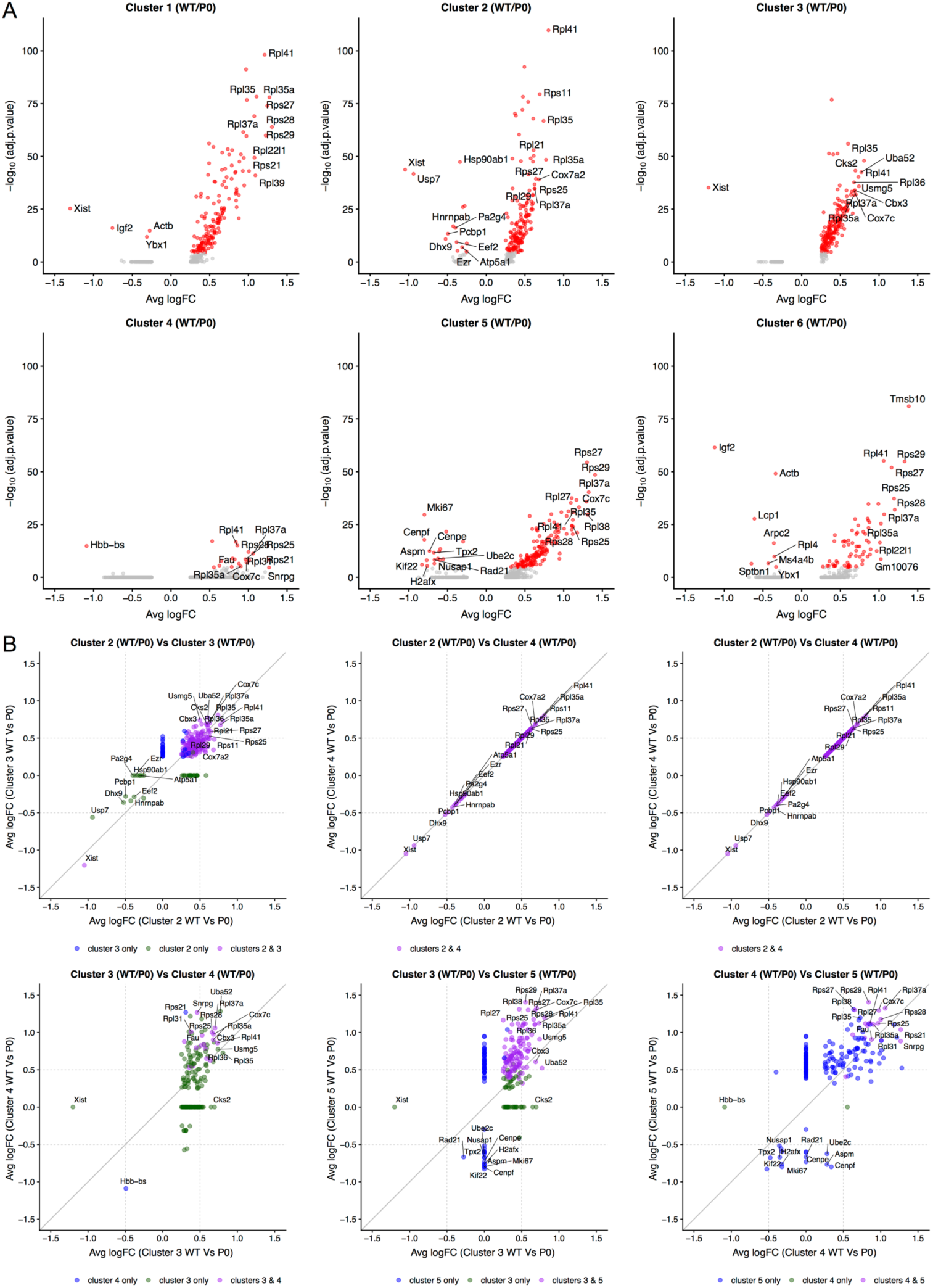
Differential transcript levels between WT and P0 cells within the six largest clusters (1-6). (**A**) Volcano plots for differential transcript levels for WT vs P0 genotype cells within each cluster revealing the ribosomal protein genes as the dominant set of genes. The top 10 genes are labelled for +/- average log fold change. (**B**) Comparison of the transcript level fold change between DN / DP and DP / T_Mat_ cell type clusters shows a similar enrichment for the ribosomal protein genes (purple).

**Figure S9:**
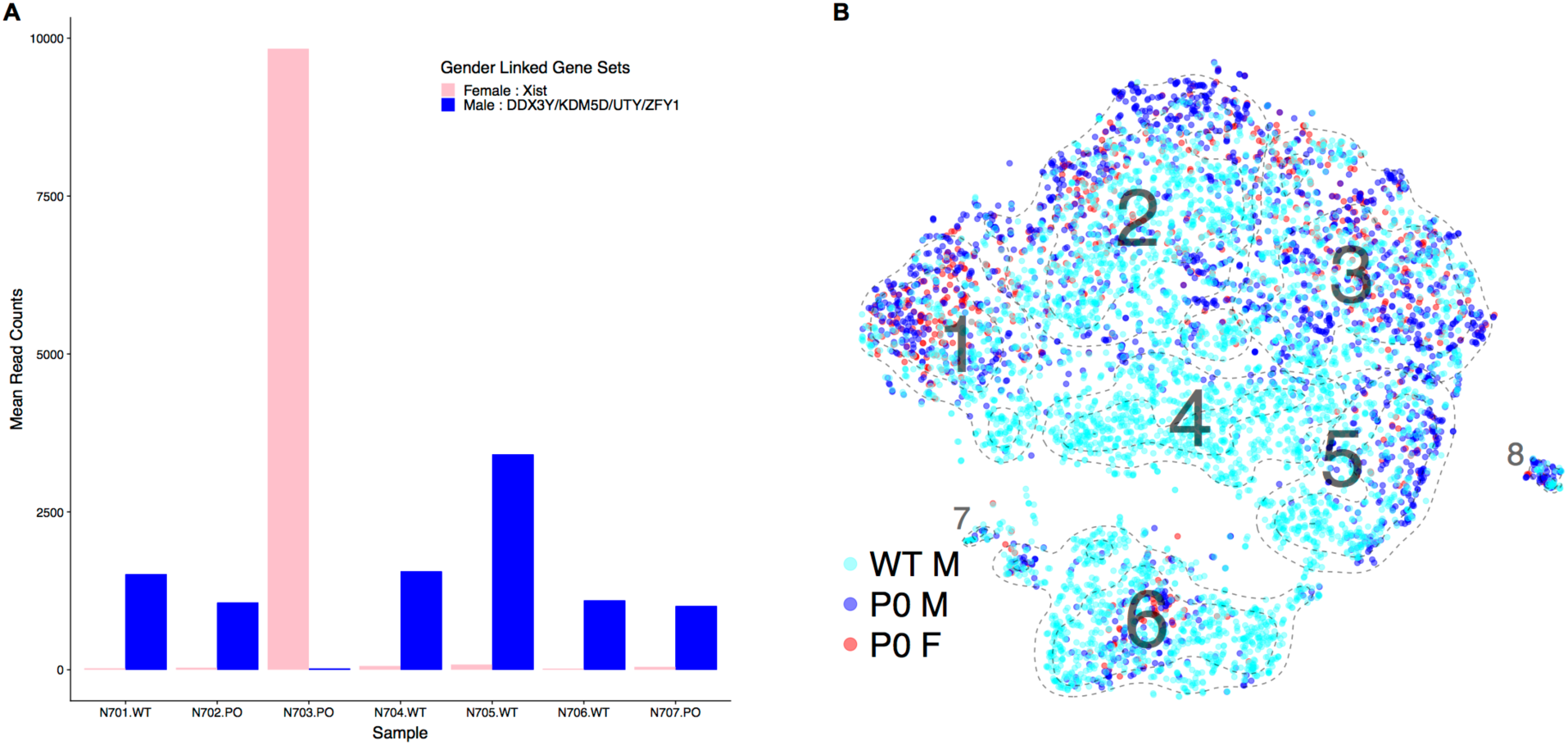
Cell clustering by sample gender. (**A**) Gender linked gene transcript levels are used to assign the gender to each of the seven samples (4 WT, 3 P_0_). (**B**) Each cell is colored by its genotype and assigned gender in the tSNE revealing no obvious gender bias within clusters.

**Table S1.** Antibodies Used in Flow Cytometry.

| **Marker** | **Clone** | **Company** |
| --- | --- | --- |
| CD11b | M1/70 | BD Biosciences |
| CD11c | N418 | BioLegend |
| CD19 | 1D3 | BD Biosciences |
| CD25 | PC61 | BioLegend |
| CD3 | 17A2, | BioLegend |
| CD4 | RM4-5 | eBioscience |
| CD44 | IM7, | BioLegend |
| CD8a | 53-6.7 | eBioscience |
| Eomes | Dan11mag | eBioscience |
| F4/80 | BM8 | eBioscience |
| FOX3Pm | NRRF-30 | eBioscience |
| Gr1 | RB6-8C5 | BioLegend |
| LyC6 | AL-21 | BD Pharmingen |
| Ly6g | 1A8 | BioLegend |
| MHC-II | M5/114.15.2 | BioLegend |
| mPDCA | eBio129c | eBioscience |
| NK1.1 | PK-136 | BioLegend |
| NKp46 | 29A1.4 | BioLegend |
| SiglecF | E50-2440 | BD Bioscience |
| TCRβ | H57-597 | eBioscience |
| TCRδ | GL3 | BioLegend |
| TER-119 | TER-119 | eBioscience |

**Table S2.** Cell marker combinations used to identify splenic and thymic subsets as shown in Figures 1, 4, and 5. Pre-processing and alignment using DropSeqTools and ctr-dropseq-tools (200 genes per cell, 3 cells pergenemin)

| Cell Type | Identification |
| --- | --- |
| B cells | CD3<sup>-</sup> TER119<sup>-</sup> CD19<sup>+</sup> |
| ILC1 | CD3<sup>-</sup> TER119<sup>-</sup> CD19<sup>-</sup> NK1.1<sup>+</sup> NKp46<sup>+</sup> EOMES<sup>-</sup> |
| NK Cells | CD3<sup>-</sup> TER119<sup>-</sup> CD19<sup>-</sup> NK1.1<sup>+</sup> NKp46<sup>+</sup> EOMES<sup>+</sup> |
| Conventional | CD3<sup>-</sup> TER119<sup>-</sup> CD19<sup>-</sup> NK1.1<sup>-</sup> MHC-II high CD11c high |
| Eosinophils | CD3<sup>-</sup> TER119<sup>-</sup> CD19<sup>-</sup> NK1.1<sup>-</sup> SiglecF<sup>+</sup> SSC high |
| Monocytes | CD3<sup>-</sup> TER119<sup>-</sup> CD19<sup>-</sup> NK1.1<sup>-</sup> Ly6G<sup>-</sup> CD11b<sup>+</sup> Ly6c<sup>+</sup> |
| Neutrophils | CD3<sup>-</sup> TER119<sup>-</sup> CD19<sup>-</sup> NK1.1<sup>-</sup> Ly6G<sup>+</sup> |
| Plasmacytoid | CD3<sup>-</sup> TER119<sup>-</sup> CD19<sup>-</sup> NK1.1<sup>-</sup> mPDCA<sup>+</sup> Ly6c<sup>+</sup> |
| Macrophages | F4/80<sup>+</sup> |
| T-Cell Subtypes |  |
| Regulatory | CD19<sup>-</sup> TER119<sup>-</sup> CD3<sup>+</sup> TCRδ<sup>-</sup> TCRβ<sup>+</sup> CD4<sup>+</sup> FoxP3<sup>+</sup> |
| δγ | CD19<sup>-</sup> TER119<sup>-</sup> CD3<sup>+</sup> TCRδ<sup>+</sup> |
| αβ | CD19<sup>-</sup> TER119<sup>-</sup> CD3<sup>+</sup> TCRδ<sup>-</sup> TCRβ<sup>+</sup> |
| SP CD4<sup>+</sup> | CD11b<sup>-</sup> NK1.1<sup>-</sup> CD19<sup>-</sup> TER119<sup>-</sup> CD11c<sup>-</sup> Gr1<sup>-</sup> TCRδ<sup>-</sup> CD4<sup>+</sup> CD8<sup>-</sup> |
| SP CD8<sup>+</sup> | CD11b<sup>-</sup> NK1.1<sup>-</sup> CD19<sup>-</sup> TER119<sup>-</sup> CD11c<sup>-</sup> Gr1<sup>-</sup> TCRδ<sup>-</sup> CD4<sup>-</sup> CD8<sup>+</sup> |
| DN | CD11b<sup>-</sup> NK1.1<sup>-</sup> CD19<sup>-</sup> TER119<sup>-</sup> CD11c<sup>-</sup> Gr1<sup>-</sup> TCRδ<sup>-</sup> |
| DP | CD11b<sup>-</sup> NK1.1<sup>-</sup> CD19<sup>-</sup> TER119<sup>-</sup> CD11c<sup>-</sup> Gr1<sup>-</sup> TCRδ<sup>-</sup> CD4<sup>+</sup> CD8<sup>+</sup> |
| DN1 | CD11b<sup>-</sup> NK1.1<sup>-</sup> CD19<sup>-</sup> TER119<sup>-</sup> CD11c<sup>-</sup> Gr1<sup>-</sup> TCRδ<sup>-</sup> |
| DN2 | CD11b<sup>-</sup> NK1.1<sup>-</sup> CD19<sup>-</sup> TER119<sup>-</sup> CD11c<sup>-</sup> Gr1<sup>-</sup> TCRδ<sup>-</sup> |
| DN3 | CD11b<sup>-</sup> NK1.1<sup>-</sup> CD19<sup>-</sup> TER119<sup>-</sup> CD11c<sup>-</sup> Gr1<sup>-</sup> TCRδ<sup>-</sup> |
| DN4 | CD11b<sup>-</sup> NK1.1<sup>-</sup> CD19<sup>-</sup> TER119<sup>-</sup> CD11c<sup>-</sup> Gr1<sup>-</sup> TCRδ<sup>-</sup> |

**Table S3.**
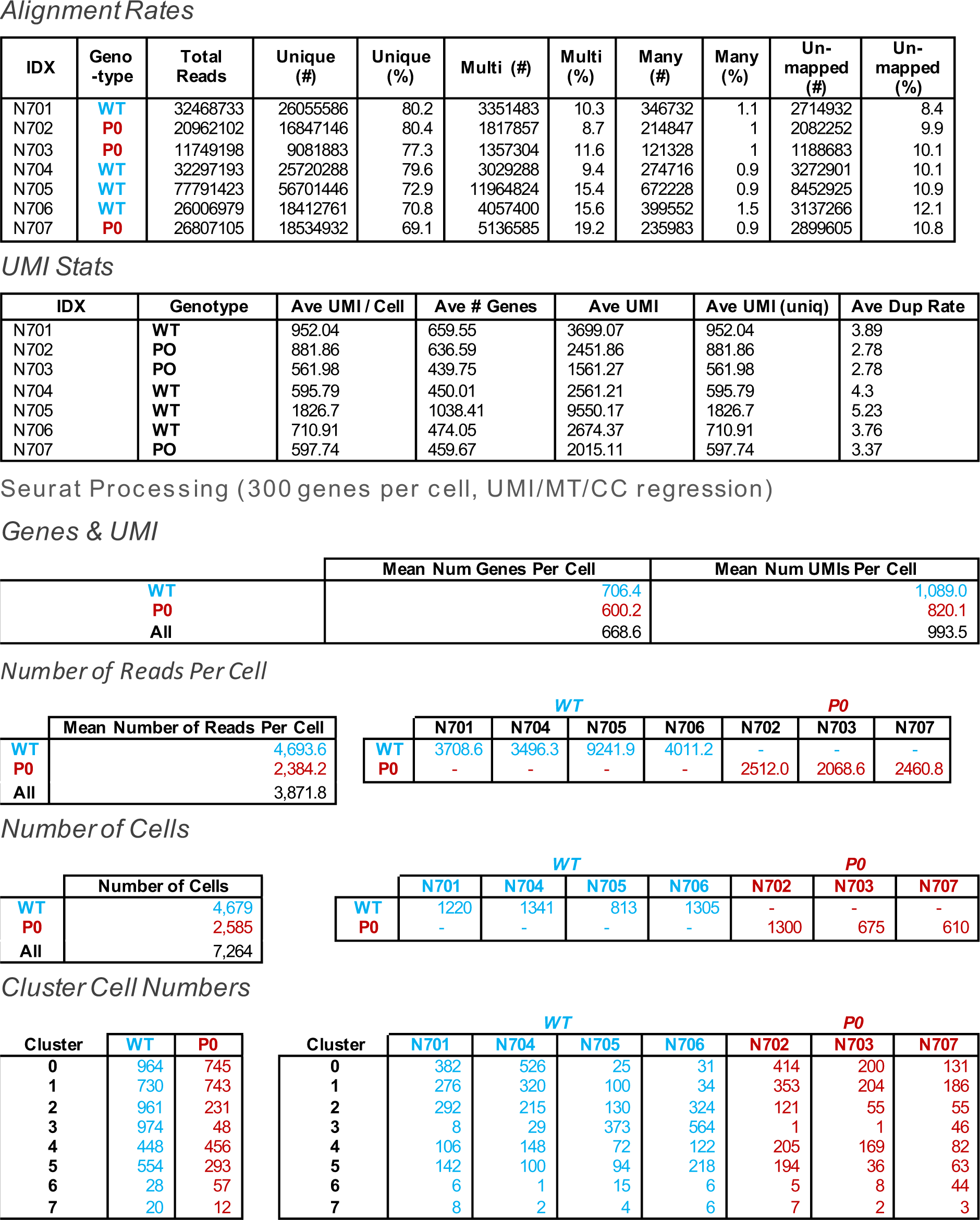
Sequencing and processing statistics for single-cell sequencing.

| IDX | Geno-type | Total Reads | Unique (#) | Unique (%) | Multi (#) | Multi (%) | Many (#) | Many (%) | Un-mapped (#) | Un-mapped (%) |
| --- | --- | --- | --- | --- | --- | --- | --- | --- | --- | --- |
| N701 | WT | 32468733 | 26055586 | 80.2 | 3351483 | 10.3 | 346732 | 1.1 | 2714932 | 8.4 |
| N702 | P0 | 20962102 | 16847146 | 80.4 | 1817857 | 8.7 | 214847 | 1 | 2082252 | 9.9 |
| N703 | P0 | 11749198 | 9081883 | 77.3 | 1357304 | 11.6 | 121328 | 1 | 1188683 | 10.1 |
| N704 | WT | 32297193 | 25720288 | 79.6 | 3029288 | 9.4 | 274716 | 0.9 | 3272901 | 10.1 |
| N705 | WT | 77791423 | 56701446 | 72.9 | 11964824 | 15.4 | 672228 | 0.9 | 8452925 | 10.9 |
| N706 | WT | 26006979 | 18412761 | 70.8 | 4057400 | 15.6 | 399552 | 1.5 | 3137266 | 12.1 |
| N707 | P0 | 26807105 | 18534932 | 69.1 | 5136585 | 19.2 | 235983 | 0.9 | 2899605 | 10.8 |

*UMI Stats*
| IDX | Genotype | Ave UMI / Cell | Ave # Genes | Ave UMI | Ave UMI (uniq) | Ave Dup Rate |
| --- | --- | --- | --- | --- | --- | --- |
| N701 | WT | 952.04 | 659.55 | 3699.07 | 952.04 | 3.89 |
| N702 | PO | 881.86 | 636.59 | 2451.86 | 881.86 | 2.78 |
| N703 | PO | 561.98 | 439.75 | 1561.27 | 561.98 | 2.78 |
| N704 | WT | 595.79 | 450.01 | 2561.21 | 595.79 | 4.3 |
| N705 | WT | 1826.7 | 1038.41 | 9550.17 | 1826.7 | 5.23 |
| N706 | WT | 710.91 | 474.05 | 2674.37 | 710.91 | 3.76 |
| N707 | PO | 597.74 | 459.67 | 2015.11 | 597.74 | 3.37 |

Seurat Processing (300 genes per cell, UMI/MT/CC regression) *Genes & UMI*
|  | Mean Num Genes Per Cell | Mean Num UMIs Per Cell |
| --- | --- | --- |
| WT | 706.4 | 1,089.0 |
| P0 | 600.2 | 820.1 |
| All | 668.6 | 993.5 |

*Number of Reads Per Cell*
| Mean Number of Reads Per Cell |  | WT |  |  |  | P0 |  |  |
| --- | --- | --- | --- | --- | --- | --- | --- | --- |
|  |  | N701 | N704 | N705 | N706 | N702 | N703 | N707 |
| WT | 4,693.6 | 3708.6 | 3496.3 | 9241.9 | 4011.2 | - | - | - |
| P0 | 2,384.2 | - | - | - | - | 2512.0 | 2068.6 | 2460.8 |
| All | 3,871.8 |  |  |  |  |  |  |  |

*Number of Cells*
| Number of Cells |  | WT |  |  |  | P0 |  |  |
| --- | --- | --- | --- | --- | --- | --- | --- | --- |
|  |  | N701 | N704 | N705 | N706 | N702 | N703 | N707 |
| WT | 4,679 | 1220 | 1341 | 813 | 1305 | - | - | - |
| P0 | 2,585 | - | - | - | - | 1300 | 675 | 610 |
| All | 7,264 |  |  |  |  |  |  |  |

|  |  |  | WT |  |  |  | P0 |  |  |
| --- | --- | --- | --- | --- | --- | --- | --- | --- | --- |
| Cluster | WT | P0 | N701 | N704 | N705 | N706 | N702 | N703 | N707 |
| 0 | 964 | 745 | 382 | 526 | 25 | 31 | 414 | 200 | 131 |
| 1 | 730 | 743 | 276 | 320 | 100 | 34 | 353 | 204 | 186 |
| 2 | 961 | 231 | 292 | 215 | 130 | 324 | 121 | 55 | 55 |
| 3 | 974 | 48 | 8 | 29 | 373 | 564 | 1 | 1 | 46 |
| 4 | 448 | 456 | 106 | 148 | 72 | 122 | 205 | 169 | 82 |
| 5 | 554 | 293 | 142 | 100 | 94 | 218 | 194 | 36 | 63 |
| 6 | 28 | 57 | 6 | 1 | 15 | 6 | 5 | 8 | 44 |
| 7 | 20 | 12 | 8 | 2 | 4 | 6 | 7 | 2 | 3 |

## References

1. Romo A, Carceller R, Tobajas J. Intrauterine growth retardation (IUGR): epidemiology and etiology. Pediatr Endocrinol Rev (2009) 6 Suppl 3:332–336.

2. Tröger B, Müller T, Faust K, Bendiks M, Bohlmann MK, Thonnissen S, Herting E, Göpel W, Härtel C. Intrauterine growth restriction and the innate immune system in preterm infants of ≤32 weeks gestation. Neonatology (2013) 103:199–204.

3. Sharma D, Shastri S, Sharma P. Intrauterine Growth Restriction: Antenatal and Postnatal Aspects. Clin Med Insights Pediatr (2016) 10:67–83.

4. Lumey LH, Stein AD, Kahn HS, van der Pal-de Bruin KM, Blauw GJ, Zybert PA, Susser ES. Cohort profile: the Dutch Hunger Winter families study. Int J Epidemiol (2007) 36:1196–1204.

5. Gaccioli F, Lager S. Placental Nutrient Transport and Intrauterine Growth Restriction. Front Physiol (2016) 7: doi:10.3389/fphys.2016.00040

6. Sferruzzi-Perri AN, Sandovici I, Constancia M, Fowden AL. Placental phenotype and the insulin-like growth factors: resource allocation to fetal growth. J Physiol (2017) 595:5057–5093.

7. Karlberg J, Albertsson-Wikland K. Growth in Full-Term Small-for-Gestational-Age Infants: From Birth to Final Height. Pediatr Res (1995) 38:733–739.

8. Salam RA, Das JK, Bhutta ZA. Impact of intrauterine growth restriction on long-term health. Curr Opin Clin Nutr Metab Care (2014) 17:249–254.

9. Melo AS, Vieira CS, Barbieri MA, Rosa-E-Silva ACJS, Silva AAM, Cardoso VC, Reis RM, Ferriani RA, Silva-de-Sá MF, Bettiol H. High prevalence of polycystic ovary syndrome in women born small for gestational age. Hum Reprod (2010) 25:2124–2131.

10. Lund LK, Vik T, Lydersen S, Løhaugen GCC, Skranes J, Brubakk A-M, Indredavik MS. Mental health, quality of life and social relations in young adults born with low birth weight. Health Qual Life Outcomes (2012) 10:146.

11. Howe LD, Chaturvedi N, Lawlor DA, Ferreira DLS, Fraser A, Davey Smith G, Tilling K, Hughes AD. Rapid increases in infant adiposity and overweight/obesity in childhood are associated with higher central and brachial blood pressure in early adulthood. J Hypertens (2014) 32:1789–1796.

12. Calkins K, Devaskar SU. Fetal origins of adult disease. Curr Probl Pediatr Adolesc Health Care (2011) 41:158–176.

13. Heindel JJ, Balbus J, Birnbaum L, Brune-Drisse MN, Grandjean P, Gray K, Landrigan PJ, Sly PD, Suk W, Cory Slechta D, et al Developmental Origins of Health and Disease: Integrating Environmental Influences. Endocrinology (2015) 156:3416–3421.

14. Bhasin KKS, van Nas A, Martin LJ, Davis RC, Devaskar SU, Lusis AJ. Maternal Low-Protein Diet or Hypercholesterolemia Reduces Circulating Essential Amino Acids and Leads to Intrauterine Growth Restriction. Diabetes (2008) 58:559–566.

15. Wang KCW, Larcombe AN, Berry LJ, Morton JS, Davidge ST, James AL, Noble PB. Foetal growth restriction in mice modifies postnatal airway responsiveness in an age and sex-dependent manner. Clin Sci (2017) 132:273–284.

16. Constância M, Hemberger M, Hughes J, Dean W, Ferguson-Smith A, Fundele R, Stewart F, Kelsey G, Fowden A, Sibley C, et al Placental-specific IGF-II is a major modulator of placental and fetal growth. Nature (2002) 417:945–948.

17. Dilworth MR, Kusinski LC, Cowley E, Ward BS, Husain SM, Constância M, Sibley CP, Glazier JD. Placental-specific Igf2 knockout mice exhibit hypocalcemia and adaptive changes in placental calcium transport. Proc Natl Acad Sci U S A (2010) 107:3894–3899.

18. Ergaz Z, Avgil M, Ornoy A. Intrauterine growth restriction-etiology and consequences: what do we know about the human situation and experimental animal models? Reprod Toxicol (2005) 20:301–322.

19. Mikaelsson MA, Constância M, Dent CL, Wilkinson LS, Humby T. Placental programming of anxiety in adulthood revealed by Igf2-null models. Nat Commun (2013) 4:2311.

20. Ursini G, Punzi G, Chen Q, Marenco S, Robinson JF, Porcelli A, Hamilton EG, Mitjans M, Maddalena G, Begemann M, et al Convergence of placenta biology and genetic risk for schizophrenia. Nat Med (2018) doi:10.1038/s41591-018-0021-y

21. Luyckx VA, Bertram JF, Brenner BM, Fall C, Hoy WE, Ozanne SE, Vikse BE. Effect of fetal and child health on kidney development and long-term risk of hypertension and kidney disease. Lancet (2013) 382:273–283.

22. Nicholl RM, Deenmamode JM, Gamsu HR. Intrauterine growth restriction, visceral blood flow velocity and exocrine pancreatic function. BMC Res Notes (2008) 1:115.

23. D’Inca R, Guen CG-L, Che L, Sangild PT, Le Huërou-Luron I. Intrauterine Growth Restriction Delays Feeding-Induced Gut Adaptation in Term Newborn Pigs. Neonatology (2011) 99:208–216.

24. Poulin JF, Viswanathan MN, Harris JM, Komanduri KV, Wieder E, Ringuette N, Jenkins M, McCune JM, Sékaly RP. Direct evidence for thymic function in adult humans. J Exp Med (1999) 190:479–486.

25. Prentice AM, Moore SE. Early programming of adult diseases in resource poor countries. Arch Dis Child (2005) 90:429–432.

26. Moore SE, Jalil F, Ashraf R, Szu SC, Prentice AM, Hanson LÅ. Birth weight predicts response to vaccination in adults born in an urban slum in Lahore, Pakistan. Am J Clin Nutr (2004) 80:453–459.

27. Moore SE, Cole TJ, Poskitt EM, Sonko BJ, Whitehead RG, McGregor IA, Prentice AM. Season of birth predicts mortality in rural Gambia. Nature (1997) 388:434.

28. Shah DK, Zúñiga-Pflücker JC. An overview of the intrathymic intricacies of T cell development. J Immunol (2014) 192:4017–4023.

29. Fischer A, de Saint Basile G, Le Deist F. CD3 deficiencies. Curr Opin Allergy Clin Immunol (2005) 5:491–495.

30. Buckner JH. Mechanisms of impaired regulation by CD4(+)CD25(+)FOXP3(+) regulatory T cells in human autoimmune diseases. Nat Rev Immunol (2010) 10:849–859.

31. Macosko EZ, Basu A, Satija R, Nemesh J, Shekhar K, Goldman M, Tirosh I, Bialas AR, Kamitaki N, Martersteck EM, et al Highly Parallel Genome-wide Expression Profiling of Individual Cells Using Nanoliter Droplets. Cell (2015) 161:1202–1214.

32. Svensson V, Vento-Tormo R, Teichmann SA. Exponential scaling of single-cell RNA-seq in the past decade. Nat Protoc (2018) 13:599–604.

33. Villani A-C, Shekhar K. Single-Cell RNA Sequencing of Human T Cells. Methods Mol Biol (2017) 1514:203–239.

34. Mahata B, Zhang X, Kolodziejczyk AA, Proserpio V, Haim-Vilmovsky L, Taylor AE, Hebenstreit D, Dingler FA, Moignard V, Göttgens B, et al Single-cell RNA sequencing reveals T helper cells synthesizing steroids de novo to contribute to immune homeostasis. Cell Rep (2014) 7:1130–1142.

35. Constância M, Hemberger M, Hughes J, Dean W, Ferguson-Smith A, Fundele R, Stewart F, Kelsey G, Fowden A, Sibley C, et al Placental-specific IGF-II is a major modulator of placental and fetal growth. Nature (2002) 417:945–948.

36. Macosko EZ, Basu A, Satija R, Nemesh J, Shekhar K, Goldman M, Tirosh I, Bialas AR, Kamitaki N, Martersteck EM, et al Highly Parallel Genome-wide Expression Profiling of Individual Cells Using Nanoliter Droplets. Cell (2015) 161:1202–1214.

37. Butler A, Hoffman P, Smibert P, Papalexi E, Satija R. Integrating single-cell transcriptomic data across different conditions, technologies, and species. Nat Biotechnol (2018) 36:411–420.

38. Lun ATL, McCarthy DJ, Marioni JC. A step-by-step workflow for low-level analysis of single-cell RNA-seq data with Bioconductor. F1000Res (2016) 5:2122.

39. Zappia L, Oshlack A. Clustering trees: a visualisation for evaluating clusterings at multiple resolutions. Gigascience (2018) doi:10.1093/gigascience/giy083

40. Moore T, Constancia M, Zubair M, Bailleul B, Feil R, Sasaki H, Reik W. Multiple imprinted sense and antisense transcripts, differential methylation and tandem repeats in a putative imprinting control region upstream of mouse Igf2. Proc Natl Acad Sci U S A (1997) 94:12509–12514.

41. Zhang MJ, Ntranos V, Tse D. One read per cell per gene is optimal for single-cell RNA-Seq. (2018) doi:10.1101/389296

42. Proserpio V, Piccolo A, Haim-Vilmovsky L, Kar G, Lönnberg T, Svensson V, Pramanik J, Natarajan KN, Zhai W, Zhang X, et al Single-cell analysis of CD4+ T-cell differentiation reveals three major cell states and progressive acceleration of proliferation. Genome Biol (2016) 17:103.

43. Dik WA, Pike-Overzet K, Weerkamp F, de Ridder D, de Haas EFE, Baert MRM, van der Spek P, Koster EEL, Reinders MJT, van Dongen JJM, et al New insights on human T cell development by quantitative T cell receptor gene rearrangement studies and gene expression profiling. J Exp Med (2005) 201:1715–1723.

44. Kisielow J, Nairn AC, Karjalainen K. TARPP, a novel protein that accompanies TCR gene rearrangement and thymocyte education. Eur J Immunol (2001) 31:1141–1149.

45. Shi Z, Fujii K, Kovary KM, Genuth NR, Röst HL, Teruel MN, Barna M. Heterogeneous Ribosomes Preferentially Translate Distinct Subpools of mRNAs Genome-wide. Mol Cell (2017) 67:71–83.e7.

46. Gui J, Mustachio LM, Su D-M, Craig RW. Thymus Size and Age-related Thymic Involution: Early Programming, Sexual Dimorphism, Progenitors and Stroma. Aging Dis (2012) 3:280–290.

47. Markert ML, Alexieff MJ, Li J, Sarzotti M, Ozaki DA, Devlin BH, Sempowski GD, Rhein ME, Szabolcs P, Hale LP, et al Complete DiGeorge syndrome: development of rash, lymphadenopathy, and oligoclonal T cells in 5 cases. J Allergy Clin Immunol (2004) 113:734–741.

48. Itoh S, Ohno T, Kakizaki S, Ichinohasama R. Epstein-Barr virus-positive T-cell lymphoma cells having chromosome 22q11.2 deletion: an autopsy report of DiGeorge syndrome. Hum Pathol (2011) 42:2037–2041.

49. Berends LM, Fernandez-Twinn DS, Martin-Gronert MS, Cripps RL, Ozanne SE. Catch-up growth following intra-uterine growth-restriction programmes an insulin-resistant phenotype in adipose tissue. Int J Obes (2012) 37:1051–1057.

50. Kernfeld EM, Genga RMJ, Neherin K, Magaletta ME, Xu P, Maehr R. A Single-Cell Transcriptomic Atlas of Thymus Organogenesis Resolves Cell Types and Developmental Maturation. Immunity (2018) doi:10.1016/j.immuni.2018.04.015

51. Sanchez CG, Teixeira FK, Czech B, Preall JB, Zamparini AL, Seifert JRK, Malone CD, Hannon GJ, Lehmann R. Regulation of Ribosome Biogenesis and Protein Synthesis Controls Germline Stem Cell Differentiation. Cell Stem Cell (2016) 18:276–290.

52. Boria I, Garelli E, Gazda HT, Aspesi A, Quarello P, Pavesi E, Ferrante D, Meerpohl JJ, Kartal M, Da Costa L, et al The ribosomal basis of Diamond-Blackfan Anemia: mutation and database update. Hum Mutat (2010) 31:1269–1279.

53. Zhou C, Zang D, Jin Y, Wu H, Liu Z, Du J, Zhang J. Mutation in ribosomal protein L21 underlies hereditary hypotrichosis simplex. Hum Mutat (2011) 32:710–714.

54. Li Z, Dordai DI, Lee J, Desiderio S. A Conserved Degradation Signal Regulates RAG-2 Accumulation during Cell Division and Links V(D)J Recombination to the Cell Cycle. Immunity (1996) 5:575–589.

55. Fagnoni FF, Vescovini R, Passeri G, Bologna G, Pedrazzoni M, Lavagetto G, Casti A, Franceschi C, Passeri M, Sansoni P. Shortage of circulating naive CD8(+) T cells provides new insights on immunodeficiency in aging. Blood (2000) 95:2860–2868.

56. Qi Q, Liu Y, Cheng Y, Glanville J, Zhang D, Lee J-Y, Olshen RA, Weyand CM, Boyd SD, Goronzy JJ. Diversity and clonal selection in the human T-cell repertoire. Proc Natl Acad Sci U S A (2014) 111:13139–13144.

57. Gui J, Morales AJ, Maxey SE, Bessette KA, Ratcliffe NR, Kelly JA, Craig RW. MCL1 increases primitive thymocyte viability in female mice and promotes thymic expansion into adulthood. Int Immunol (2011) 23:647–659.

